# Divergent modulation of dopaminergic neurons by hypocretin/orexin receptors-1 and -2 shapes dopaminergic cell activity and socio-emotional behavior

**DOI:** 10.1101/2025.02.14.638329

**Authors:** Stamatina Tzanoulinou, Simone Astori, Laura Clara Grandi, Francesca Gullo, Richie Kalusivikako, Simran Rai, Galina Limorenko, Mehdi Tafti, Andrea Becchetti, Anne Vassalli

## Abstract

**BACKGROUND:** Many neuropsychiatric disorders involve dysregulation of the dopaminergic (DA) input to the forebrain. Of particular relevance are DA projections stemming from the midbrain ventral tegmental area (VTA). A key neuromodulatory influence onto DA^VTA^ neurons arises from lateral hypothalamic area hypocretin/orexin (OX) neurons. Despite being a major input, the differential action of orexin peptides A and B (OXA and OXB) on orexin receptors 1 and 2 in DA cells is poorly understood. We recently identified profoundly divergent functions of OX_1_R *vs* OX_2_R in DA cells in regulating sleep/wake architecture, brain oscillations and cognitive behaviors. OX_2_R, but not OX_1_R, loss dramatically increased time in EEG theta-rich (alert) wakefulness, reward-driven learning and attentional skills, but impaired inhibitory control.

**METHODS:** Using genetically engineered mice whose DA cells selectively lack OX input via *Hcrtr1* (DA*^Ox1R-KO^*) or *Hcrtr2* (DA*^Ox2R-KO^*), we assessed intrinsic excitability and electrophysiological responses of DA^VTA^ neurons and evaluated behavioral phenotypes across multiple domains.

**RESULTS:** We uncover previously unrecognized effects of OX peptides on DA^VTA^ cell response. In WT and control mice, we show that while OXA enhances, OXB diminishes DA^VTA^ neuronal excitability. OX_1_R-deficient DA cells lose OXA responding and OX_2_R-deficient DA cells lose OXB responding. DA *Ox1R* loss generates anxiety-like behavior and context-dependent hyperactivity. In contrast, OX_2_R loss decreases sociability and, despite exhibiting enhanced reward-driven learning, mice show highly compromised aversion-driven learning.

**CONCLUSIONS:** We evidence strikingly distinct functions of OX_1_R vs OX_2_R signaling in modulating the intrinsic excitability of DA^VTA^ neurons and influencing DA-related behaviors. These data implicate OX→DA signaling pathways in neuropsychiatric endophenotypes relevant to obsessive-compulsive, attention-deficit/hyperactivity, and autism spectrum disorders, and raise important considerations for the development of OXR-targeted therapeutics.

## INTRODUCTION

Neuromodulators are small neurotransmitters or neuropeptides released by neurons that regulate cellular and synaptic properties across broad neural territories, thereby enabling coordinated reconfigurations of neural circuit output and transitions of brain states. These effects may be exerted primarily by volume transmission, targeting both synaptic and extrasynaptic receptors.

Noradrenaline (NA), dopamine (DA), serotonin (5-hydroxytryptamine, 5HT), histamine, acetylcholine, and hypocretin/orexin (OX) neuropeptides are the main known wake-promoting neuromodulators. DA and OX are also key hubs of the motivational network^1^. OX are uniquely produced by ∼80’000 hypothalamic glutamatergic cells of the lateral hypothalamus area (LHA) in humans (or about 1’000 in mice)^2,3^ that release the 2 neuropeptides hypocretin-1 and -2^4^, also called orexin A and B (OXA and OXB)^5^, central for arousal induction and maintenance. OXs are also critically implicated in energy homeostasis, drug reward, autonomic and endocrine regulation, anxiety and depression-related behaviors^6,7^.

OX pleiotropism is made possible by the brain-wide extent of OX circuitry. OX cells are multi-modal sensors and integrators of internal and external environmental signals, which broadcast effector signals through projections to all wake-promoting monoaminergic and cholinergic nuclei, as well as to their targets in the neocortex, thalamus, hippocampus, amygdala and spinal cord^8^. OX cells fire maximally during active wakefulness, consistently with their role in sustaining heightened arousal, but also display transient firing during non-rapid-eye-movement sleep (NREMS)^9^ and phasic REMS^10^. Their activity during sleep was recently conceptualized as critical for OX role in stabilizing vigilance states^11^ and controlling timely sleep-to-wake transitions^12,13^. OX loss severely compromises the integrity and stability of the 3 cardinal vigilance states: wakefulness, NREMS and REMS^11,14^, causing a lifelong debilitating disease, narcolepsy type-1^15^.

OXs act through two genetically independent and differentially expressed GPCRs, OX_1_R and OX_2_R, encoded by the *Hcrtr1* and *Hcrtr2* genes. OXA binds both OX_1_R and OX_2_R with high affinity, while OXB has higher affinity for OX_2_R^5,16^. Although genetic evidence in dogs and mice points to a stronger involvement of OX_2_R in vigilance state regulation, most studies focus on OXA and OX_1_R function, resulting in underexplored appreciation of the specific functions of OXB and OX_2_R signaling. Brain levels of the two peptides, and differential signaling and role of the two receptors remain largely unknown, despite their suspected role in many disorders.

OX neurons respond to salient stimuli of either negative or positive valence across multiple metabolic and sensory modalities and are implicated in stress- and reward-associated plasticity. Low glucose or O_2_, CO_2_ or pH changes, visual, acoustic, and olfactory stimuli, all activate OX neurons^7^. Positive stimuli such as highly palatable food, emotional elation, social reunion can trigger cataplexy, the decoupling of wakefulness from motor tone, NT1 pathognomonic symptom and signature of OX signaling deficiency^17^.

As mentioned, OX neurons exert their effects by projecting to brain areas controlling arousal, motivation and stress responses, including the basal forebrain, locus coeruleus (LC), dorsal raphe, extended amygdala, ventral tegmental area (VTA), nucleus accumbens (NAc) and medial prefrontal cortex (mPFC)^8,18,19^. Data implicate DA as a major OX effector. For instance, while daytime sleepiness and cataplexy both stem from OX deficiency and represent NT1 two major symptoms, they arise from fundamentally distinct mechanisms^17^, yet both respond to dopaminergic drugs. In narcoleptic dogs and mice, sleep attacks respond to D_1_R agonists and cataplexy responds to D_2/3_R antagonists^20,21^. OX cells densely project to DA^VTA^ neurons, which express both *Hcrtr1*/*Ox_1_R* and *Hcrtr2*/*Ox_2_R*^22^, and OX^LH^→DA^VTA^ connectivity is thought to be critical in the behavioral implications of the OX-DA interplay^7,23–31^. The precise role of OX at the interface of the VTA→NAc mesolimbic and VTA→PFC mesocortical systems, and OX_1_R and OX_2_R relative contributions in the modulation of reward, motivation, and executive functions are however largely unknown.

From a cellular standpoint, OX stimulation elicits different firing patterns in different DA^VTA^ cell subsets, and single DA^VTA^ cells express either *Hcrtr1*, *Hcrtr2*, or both receptors^32^. In general, the prevalent effect of OXA on DA^VTA^ cells appears excitatory, as OXA elevates [Ca^2+^]_cytosolic_ in isolated rat DA^VTA^ cells^33^, and enhances DA^VTA^ firing and induce tetrodotoxin-resistant depolarization in brain slices^32^. OX is thought to stimulate DA^VTA^ activity both through direct OXR activation at DA somatodendritic cell membranes, and indirectly by potentiating glutamatergic afferents onto DA cells^32,34,35^. Borgland *et al* showed that OXA potentiates N-methyl-D-aspartate receptor (NMDAR) transmission in DA^VTA^ synapses *ex vivo*, while OX_1_R antagonism blocks locomotor sensitization to cocaine in mice *in vivo*, as well as occludes VTA AMPA/NMDA ratio increase *ex vivo*, a proxy for plasticity normally observed following cocaine exposure^34^. Borgland *et al* subsequently showed that optical stimulation of VTA OX terminals is insufficient by itself to alter DA release in the NAc^36^. However, when optogenetic stimulation of OX→DA fibers was paired with intra-VTA electrical stimulation, DA levels in the NAc core increased significantly—an effect that was blocked by OX_1_R antagonism. These findings suggest that OXA-mediated OX_1_R signaling potentiates VTA→NAc DA transmission, likely through recruitment of VTA glutamatergic afferents^36^.

OXB action on DA^VTA^ neurons appeared even more complex, with a subset of neurons responding to OXB by alternate periods of silent hyperpolarization and firing, thus featuring burst firing^32^. Mechanisms underlying these different responses remain unknown. Once again, they may result from integration of cell-autonomous (direct) effects of OXB on OXRs on somatodendritic DA cell membranes and indirect effects mediated by OXRs localized on local GABAergic terminals modulating DA cell activity.

To functionally and cell-specifically interrogate OX modulation of DA activity, we generated mice whose DA neurons are selectively unresponsive to OX_1_R or OX_2_R signaling by DA-specific *Hcrtr1* or *Hcrtr2* gene inactivation^22^. Surprisingly, we found that DA loss of OX_2_R, but not OX_1_R, nor loss of both receptors, induced electrocortical hyperarousal, with mice spending most of their waking time in an alert, EEG theta/fast-gamma wave-enriched substate, displaying enhanced theta-gamma phase-amplitude coupling associated with higher attentional skills and learning speed. These advantages were however compromised by impulsive- and compulsive-like behavioral patterns^22^.

Here, we systematically investigate the roles of *Hcrtr1 vs Hcrtr2* on DA neuronal excitability and how DA OX_1_R vs OX_2_R loss affects socioemotional behaviors. Our data uncover profoundly distinct functions of the two receptors, and an unsuspected OX_2_R-mediated inhibition of DA^VTA^ neurons. In light of the many OX_1_R and OX_2_R-targeted therapeutics in development^37^, our findings raise questions regarding long-term therapy of disorders using these agents.

## METHODS AND MATERIALS

Detailed description is provided in the Supplement.

### Study approval

All procedures performed in Switzerland were conducted in accordance with the Swiss National Institutional Guidelines on Animal Experimentation and approved by the Swiss Cantonal Veterinary Office Committee for Animal Experimentation of Canton de Vaud. Procedures for *ex vivo* recordings of dissociated VTA cells followed the Italian law and were approved by the Italian Ministry of Health and the local Ethical Committee.

## RESULTS

### Divergent effects of OXA and OXB peptides in dissociated DA^VTA^ neurons of WT mice

Shortly after the discovery of the hypocretin/orexin neuropeptides^4,5^, Korotkova et al reported that application of OXA on rat VTA slices enhanced the spontaneous firing frequency of DA neurons. OXB had a more heterogeneous effect, including induction of phasic firing within a subset of DA^VTA^ cells^32^. To further characterize neuromodulation of DA neuronal activity by OXs, we first performed patch-clamp recordings on isolated DA cells acutely dissociated from WT VTA slices (see DA cell identifying criteria in Materials and Methods). This preparation excluded the confounding synaptic effects incurred by other VTA network cell types. To avoid perturbation of the DA intracellular medium^38^, spontaneous action currents were recorded in cell-attached condition and reported as 1-min time-bins, before, during, and after orexin peptide application. OXA (100 nM) increased DA^VTA^ spontaneous activity in approximately 70% of the cells tested, consistently with previous data in VTA slices^32,39^, with a slow recovery (**Fig. 1a**). In contrast, within 1-2 min of exposure, we found that OXB (100 nM) generally decreased DA^VTA^ cell firing frequency by 20-25%, with a relatively rapid recovery upon peptide removal (**Fig. 1b**). When the OX_2_R-selective agonist [Ala^11^,D-Leu^15^]-Orexin B (OXB-AL^40^) (200 nM) was applied, a more robust firing rate decrease was observed, with an approximately 40% decrease on average in DA^VTA^ cell firing (**Fig. 1c**).

**Figure 1.**
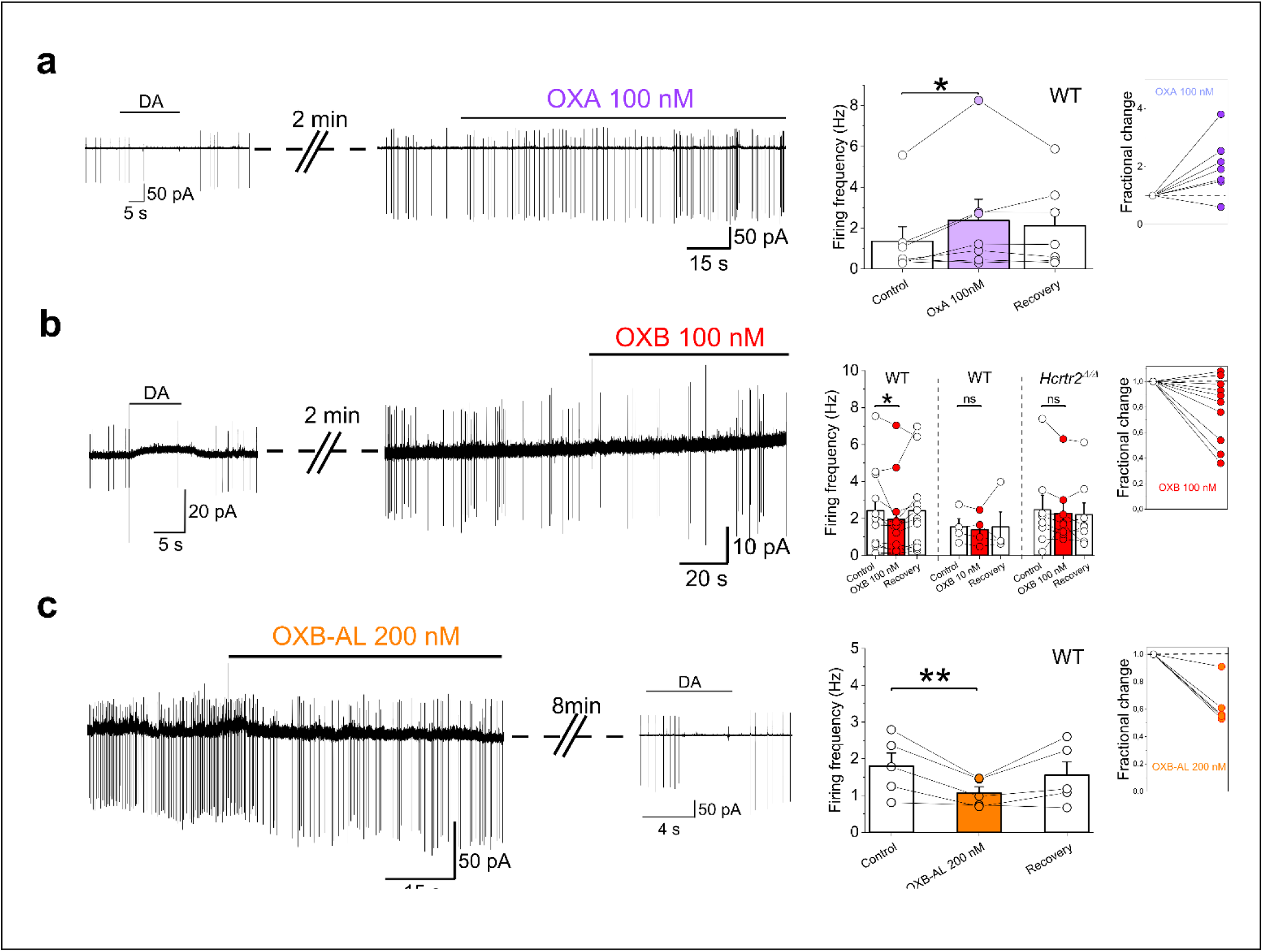
OXA increases, while OXB decreases, cell firing of dissociated DA^VTA^ cells from WT mice. DA cells were acutely isolated from VTA slices of C57BL/6J (WT) or *Hcrtr2^Δ/Δ^* (OX_2_R-deficient)^22^ mice at 2-3-weeks of age. Spontaneous action currents were assessed by patch-clamping in cell-attached mode, before, during, or after (Recovery) bath application of OXA (100 nM), OXB (10 or 100 nM), or the OX_2_R-specific agonist OXB-AL (200 nM). V_pipette_ was set to 0 mV. (**a**) (*Left*) Representative trace showing spontaneous firing of a WT DA cell exposed first to DA and next to OXA. (*Right*). Firing frequencies calculated before (Control; 2 min), during the 2^nd^ min of OXA exposure (Purple), or after washout (Recovery). Bar graphs: Control: 1.36 ± 0.717 Hz; OXA: 2.374 ± 1.05 Hz; Recovery: 2.11 ± 0.79 Hz (*χ^2^* = 6, *P* = 0.0498; Friedman ANOVA; post-hoc WNMT test between Control and OXA: *P* = 0.043, n = 7). *(Inset)* Relative effect of OXA on firing activity in individual DA cells. Datapoints (Purple) are OXA/Control firing frequency ratios for each cell. Control values are set to 1. (**b**) *(Left)*. Representative trace showing spontaneous firing of a WT DA cell exposed first to DA, and next to OXB. (*Right*). Firing frequencies calculated before (Control; 2 min), during the 1st min of OXB exposure (Red), or after washout (Recovery). Left bar graphs (WT): Control: 2.42 ± 0.63 Hz; OXB 100 nM: 1.955 ± 0.59 Hz (*χ^2^* = 8.17, *P* = 0.01685; Friedman ANOVA; post-hoc WNMT test between Control and OXB: *P* = 0.022; n = 12); Recovery: 2.42 ± 0.64 Hz. Middle bar graphs (WT): Control: 1.54 ± 0.44 Hz; OXB 10 nM: 1.39 ± 0.43 Hz (*F* = 0.09834; *P* = 0.78; repeated-measures 1-way ANOVA; n = 4); Recovery: 1.54 ± 0.81 Hz. Right bar graphs (*Hcrtr2^Δ/Δ^*) depict data recorded from DA neurons isolated from VTA slices from *Hcrtr2^Δ/Δ^* mice: Control 2.46 ± 0.79 Hz; OXB 100 nM: 2.28 ± 0.62 Hz (*χ^2^* = 3.25, *P* = 0.197; Friedman ANOVA; n = 8); Recovery: 2.195 ± 0.65 Hz. (Inset) Relative effect of OXB 100 nM (Red) on firing activity in individual WT DA cells, calculated as in (a). (**c**) (*Left*) Representative trace showing spontaneous firing of WT DA cell exposed first to OXB-AL, then to DA. (*Right*) Firing frequencies calculated before (Control, 2 min), during the 1^st^ min of OXB-AL exposure (Orange), or after washout (Recovery). Bar graphs: Control: 1.8 ± 0.36 Hz; OXB-AL: 1.07 ± 0.17 Hz (*F* = 8.641, *P* = 0.01; repeated-measures 1-way ANOVA; post-hoc Holm-Sidak test between Control and OXB-AL: *P* = 0.0035; n = 5); Recovery: 1.55 ± 0.37 Hz. (*Inset*) Relative effect of OXB-AL (Orange) on firing activity in individual WT DA cells, calculated as in (a).

As OXB poorly binds OX_1_R but binds OX_2_R with high affinity, we next assessed the OXB response of DA^VTA^ dissociated neurons from *Hcrtr2^Δ/Δ^* mice, that lack the *Hcrtr2* gene, and thus OX_2_R, in all cells^22^. OXB (100 nM) was found to only exert minor effects on DA^VTA^ cells of these mice (**Fig. 1b, right**), validating the predominantly OX_2_R-dependent effect of OXB.

While these findings confirm that OXA increases DA^VTA^ neuronal firing, they also newly reveal that OXB-mediated OX_2_R signaling can induce a transient decrease in DA^VTA^ cell firing. This suggests existence of previously unrecognized differences between OX_1_R and OX_2_R signaling in DA neurons.

Our genetically-based mutants lacking OX_1_R or OX_2_R selectively in DA cells^22^ allow to parse out the different contributions of the two receptors to DA^VTA^ cell physiology. We therefore next measured how HCRT-1 (OXA) and HCRT-2 (OXB) impact the excitability of DA^VTA^ cells, and whether OX action is altered in DA*^Ox1R-KO^* and DA*^Ox2R-KO^*mice relative to their respective controls.

### Opposite effects of OX_1_R and OX_2_R activation on DA^VTA^ cell firing *ex vivo*

To corroborate the results obtained in dissociated cells and assess the impact of the selective loss of either OX_1_R or OX_2_R in DA cells, we performed electrophysiological recordings of DA^VTA^ cells in acute slices from DA*^Ox1R-KO^* and DA*^Ox2R-KO^* mice and their respective controls (DA*^Ox1R-CT^* and DA*^Ox2R-CT^*) and assessed their responsiveness to OXA and OXB.

Our conditional gene targeting design entails replacing the 1st coding exon of the *Hcrtr* gene by a GFP-reporter activated upon lox site-recombination, acting as marker of both successful Cre-mediated recombination and activity of the endogenous *Hcrtr1* or 2 gene promoter^22^. This enabled us to estimate the percentage of ventral midbrain TH^+^ cells that express *Hcrtr1* and *Hcrtr2*, at 83.0±2.8%, and 87.2±1.5% of TH+ neurons, respectively^22^. These data suggest that in the a-p range that we assay (-2.92 to -3.88 mm from bregma), a majority of ventral midbrain DA neurons express *Hcrtr1* and *Hcrtr2* (>83% and >87%, respectively), and at least 72% (0.83 x 0.87) of ventral midbrain VTA neurons express both receptors in the mixed C57BL/6J X C57BL/6N genetic background that our mice share. Relative expression of the 2 receptors may somewhat differ in other mouse strains^41^.

First, we assessed intrinsic excitability of DA^VTA^ cells from DA*^Ox1R-KO^* and DA*^Ox1R-CT^* mice through patch-clamp recordings in DA neurons of the lateral VTA (**Fig. 2a**, see criteria for cell identification in Materials and Methods), as this region contains NAc-projecting DA cells shown to be modulated by orexins^39^. When cell firing was elicited by depolarizing current injections (**Fig. 2a-b**), no differences were observed in the firing frequency-injected current relationships between DA*^Ox1R-KO^* and DA*^Ox1R-CT^* mice (**Fig. 2c**), indicating that OX_1_R loss did not alter basal intrinsic excitability of DA^VTA^ cells. To study the impact of orexin release on intrinsic excitability, we bath-applied orexin peptides and measured the changes in firing of DA^VTA^ cells elicited by brief depolarizations of constant amplitude^39^. As OXA binds OX_1_R with high affinity^5,16^, slices from DA*^Ox1R-KO^* and DA*^Ox1R-CT^* mice were perfused with OXA (100 nM) for 5 min after a 10 min baseline (**Fig. 2d-e**). Whereas OXA increased the firing rate in control mice, it had no effect in DA*^Ox1R-KO^* mice (**Fig. 2e-f**), consistent with reports of Baimel *et al*^39^, who showed that OXA-induced DA^VTA^ firing increase is OX_1_R-dependent, and cell-autonomous (i.e., preserved if glutamatergic and GABAergic synaptic receptors are blocked). Since most of DA neurons from DA*^Ox1R-KO^* mice are expected to still express OX_2_R, the absence of effect of OXA exposure also suggests that OXA-driven OX_2_R signaling is not significantly contributing in modulating the firing rate of DA cells.

**Figure 2.**
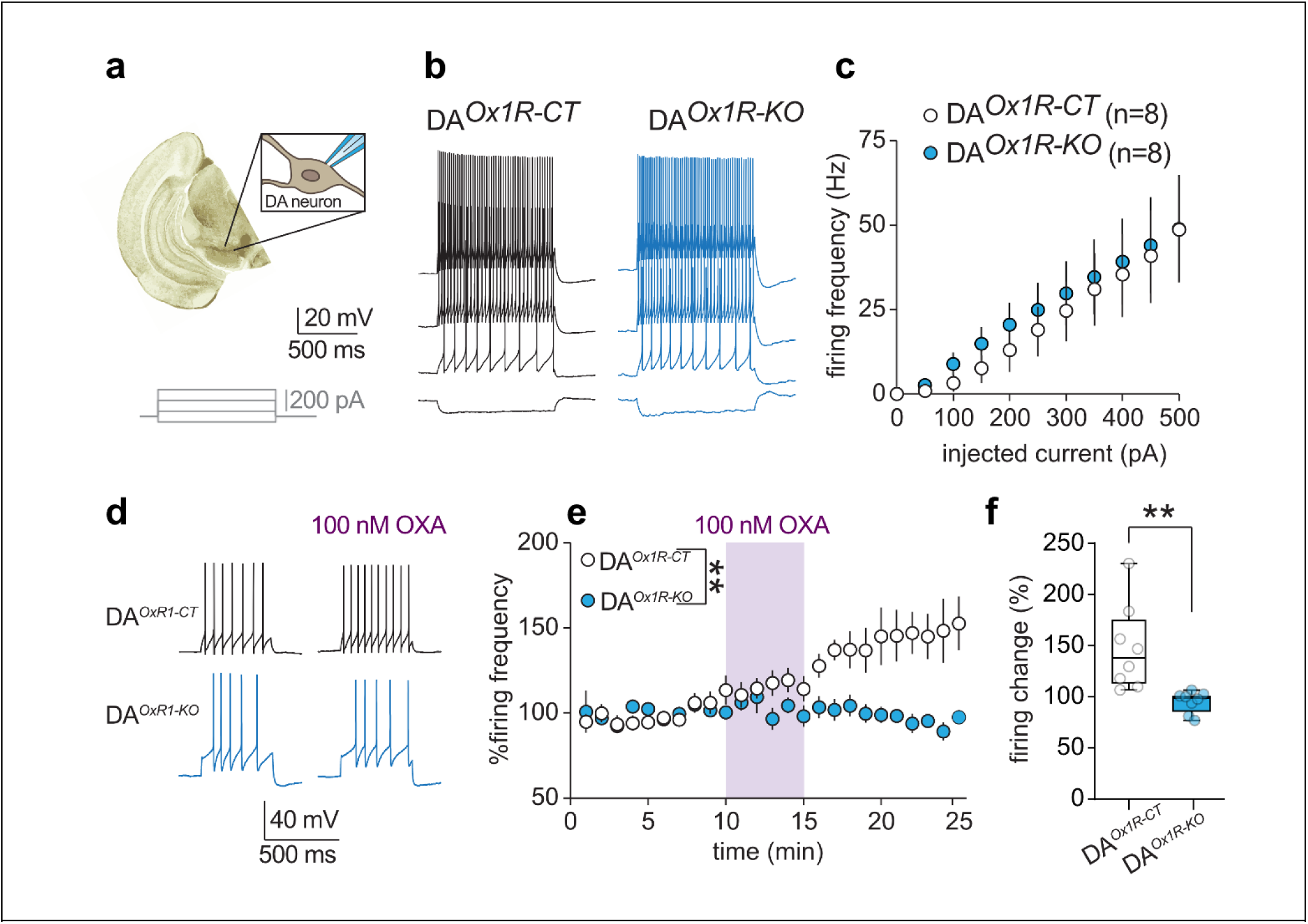
OXA enhances the excitability of DA^VTA^ neurons from control mice but not DA*^Ox1R-KO^* mice. (**a**) Scheme of *ex vivo* whole-cell patch-clamp recordings from DA neurons in VTA slices from DA*^Ox1R-CT^*and DA*^Ox1R-KO^* mice. (**b**) Representative traces of neuronal firing elicited by increasing somatic current injections. (**c**) Frequency-current relationship of action potential discharges shows no difference between CT and KO mice (2-way ANOVA for effect genotype *F*_1,14_ = 0.1426; *P =* 0.711). (**d**) Examples of neuronal firing elicited in response to a depolarizing step prior to and after bath-application of OXA (100 nM) in DA neurons from DA*^Ox1R-CT^* and DA*^Ox1R-KO^* mice. (**e**) Mean firing frequency values normalized to the first 10 min (n=8 cells per genotype). OXA increased firing in DA*^Ox1R-CT^* but not in DA*^Ox1R-KO^* mice (2-way ANOVA for effect genotype *F*_1,14_ = 10.45; *P =* 0.006 and genotype × treatment interaction *F*_24,334_ = 5.772; *P <*0.001). (**f**) Average OXA-induced change in neuronal firing in DA*^Ox1R-CT^* and DA*^Ox1R-KO^* mice (*t* (7.87) = 3.419; *P =* 0.009, *t*-test). Bar graphs show individual values and mean ± SEM. Timelines show mean ± SEM.

To examine the intrinsic excitability properties of DA^VTA^ cells lacking *Hcrtr2*, we followed a similar methodology (**Fig. 3a**) but assessed the cell response to bath-applied OXB, as OXB binds OX_2_R with high affinity but only poorly OX_1_R^5,16^. No differences in basal intrinsic excitability were observed between cells from DA*^Ox2R-KO^* and DA*^Ox2R-CT^* mice (**Fig. 3b-c**). Strikingly, and consistently with the results from dissociated VTA cells, we found that OXB decreased firing of DA^VTA^ neurons from control mice (**Fig. 3d-e-f**). This decrease was abolished in DA^VTA^ cells of DA*^Ox2R-KO^* mice, lacking an active *Hcrtr2* gene, thus validating the DA*^Ox2R-KO^* mouse model, while indicating that the OXB-induced decreased cell firing of control mice is OX_2_R-dependent. In fact, DA^VTA^ cells of DA*^Ox2R-KO^* mice showed a tendency towards increased firing upon OXB exposure (comparison between number of spikes pre- *vs* post-OXB spike in KO: *P*=0.077, **Fig. 3e**). This may reflect preserved OX_1_R signaling in DA*^Ox2R-KO^* mice activated by OXB, despite the reported lower binding affinity of OXB for OX_1_R. Importantly, these data align with those described above in acutely dissociated DA^VTA^ cells and indicate that OXB has an inhibitory action on lateral DA^VTA^ cell excitability, which is opposite to the action of OXA.

**Figure 3.**
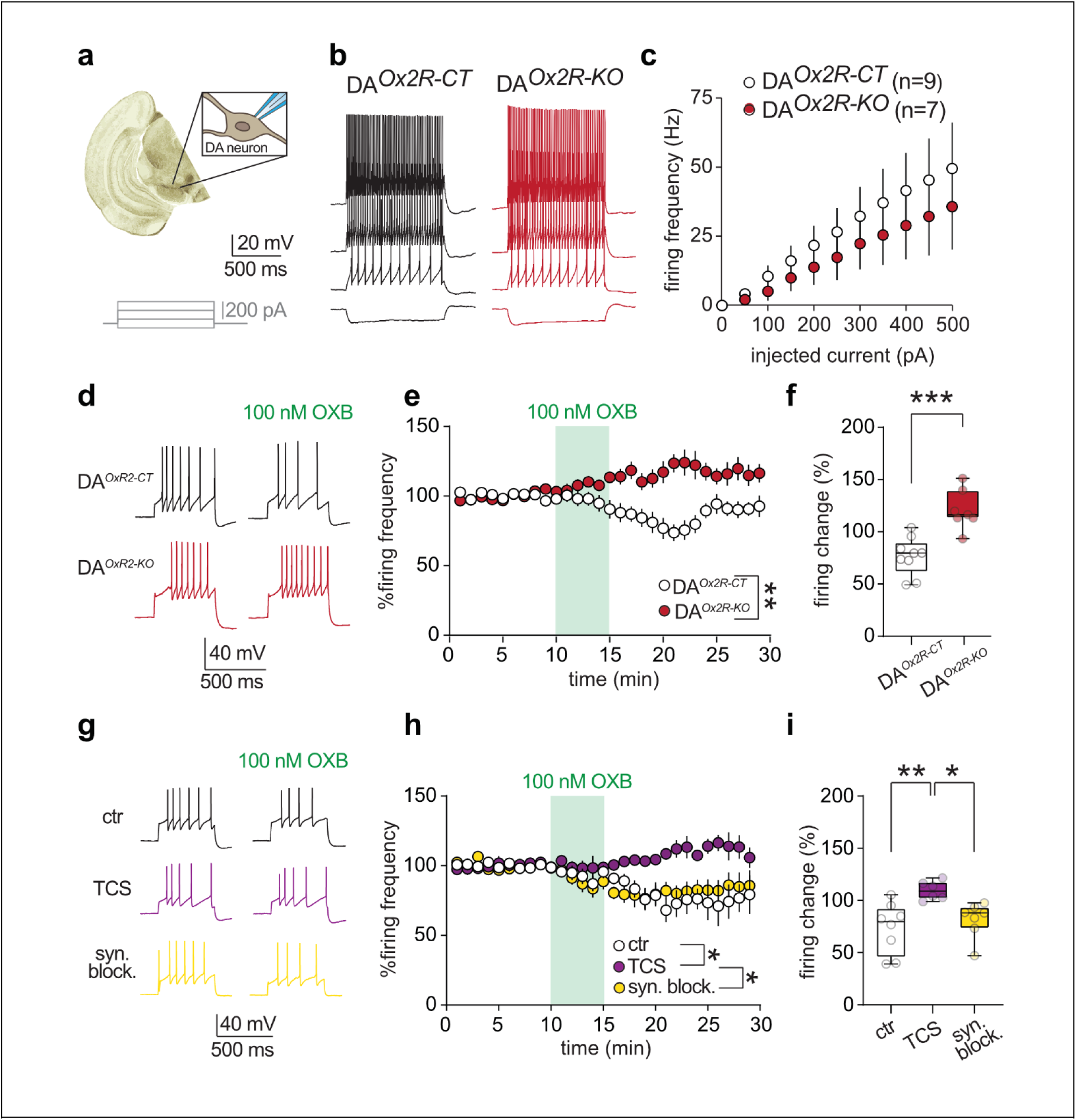
OXB decreases the excitability of DA^VTA^ neurons from control mice but not DA*^Ox2R-KO^* mice. (**a**) Scheme of *ex vivo* patch-clamp recordings from DA neurons of DA*^Ox2R-CT^* and DA*^Ox2R-KO^* mice. (**b**) Representative traces of neuronal firing elicited by increasing somatic current injections. (**c**) Frequency-current relationship of action potential discharges shows no statistical difference between CT and KO mice (2-way ANOVA for effect genotype *F*_1,14_ = 0.457; *P =* 0.510). (**d**) Examples of neuronal firing elicited in response to a depolarizing step prior to and after bath-application of OXB (100 nM) in DA neurons from DA*^Ox2R-CT^* and DA*^Ox2R-KO^* mice. (**e**) Mean firing frequency values normalized to the first 10 min (n=9:7 cells, CT:KO). OXB transiently decreased firing in DA*^Ox2R-CT^* (paired *t* test; t_8_ = 3.546; *P =* 0.008), but not DA*^Ox2R-KO^* mice, which showed instead a tendency towards increased firing (*t*_6_ = 2.132; *P =* 0.077, paired t-test). 2-way ANOVA revealed a genotype effect (*F*_1,14_ = 14.87; *P =* 0.002) and a significant genotype × treatment interaction (*F*_28,383_ = 6.484; *P <*0.001). (**f**) Average OXB-induced change in neuronal firing in DA*^Ox2R-CT^*and DA*^Ox2R-KO^* mice (*t*_12.72_ = 4.729; *P <* 0.001, *t*-test). (**g**) Representative examples of neuronal firing elicited by a depolarizing step before and after bath application of OXB (100 nM) in DA neurons from VTA slices prepared from C57BL/6J mice under three conditions: control (ctr, without blockers), in the presence of the OX_2_R antagonist TCS-OX2-29 (10 µM), and in the presence of synaptic blockers (syn. block.; 10 µM DNQX, 50 µM D,L-APV, 100 µM picrotoxin). (**h**) Mean firing frequency normalized to the first 10 min for each experimental group (n = 8, 6, 7, respectively). OXB decreased firing in the control condition (*W* = -32; *P =* 0.023, Wilcoxon test) and in the presence of synaptic blockers (*t* (6) = 2.634; *P =* 0.039, paired *t*-test), but not when OX_2_R receptors were blocked (*t*_6_ = 1.662; *P =* 0.148, paired *t*-test). A mixed-effects analysis revealed a significant effect of OXB (*F*_30, 570_ = 2.787; *P <* 0.001) and a significant effect of the blockers (*F*_2,18_ = 6.433; *P <* 0.01). (**i**) Average OXB-induced change in neuronal firing across the three groups (*F*_2,18_ = 6.815; *P =* 0.006, one-way ANOVA). Asterisks indicate statistical significance as determined by Holm-Šídák’s multiple comparisons test (*P<0.05; **P<0.01).

To corroborate this finding and further confirm that the OXB effect is cell-autonomous in VTA slices and OX_2_R-dependent, we performed experiments in C57BL/6J WT mice. Consistent with our results in DA*^Ox2R-CT^* mice, OXB decreased the firing rate of DA^VTA^ cells from WT mice. This effect was prevented by OX_2_R blockade with TCS-OX2-29 (**Fig. 3h-i**) but persisted in presence of antagonists of ionotropic synaptic receptors (**Fig. 3h-i**), ruling out possible indirect actions of OXB on DA^VTA^ cell excitability, e.g., via glutamatergic transmission^35^.

These data indicate that OXA and OXB peptides exert opposite actions on DA^VTA^ cell intrinsic excitability, and their impact is lost in our DA*^Ox1R-KO^* and DA*^Ox2R-KO^* mouse models, respectively. Having shown that our binary KO:CT models of DA *Hcrtr1* and *Hcrtr2* loss recapitulate OX_1_R and OX_2_R opposite signaling activities in DA cells of control mice, and their specific loss in DA cells of KO mice, we next aimed to determine the behavioral correlates of these two divergent OX→DA pathways by exposing the mice to a panel of tests. We selected assays representing four axes shown to heavily depend on OX and DA transmission: positive (rewarding) and negative (aversive) affect-related behaviors, anxiety-like behaviors and cognitive (attention-dependent) behaviors (**Fig. 4**). We acknowledge that this segregation is not absolute, with some degree of domain overlaps. Cognitive components for instance are also addressed by the active avoidance test used to assess negative affect. This categorization, however, offers a systematic context to present and interpret our findings.

**Figure 4.**
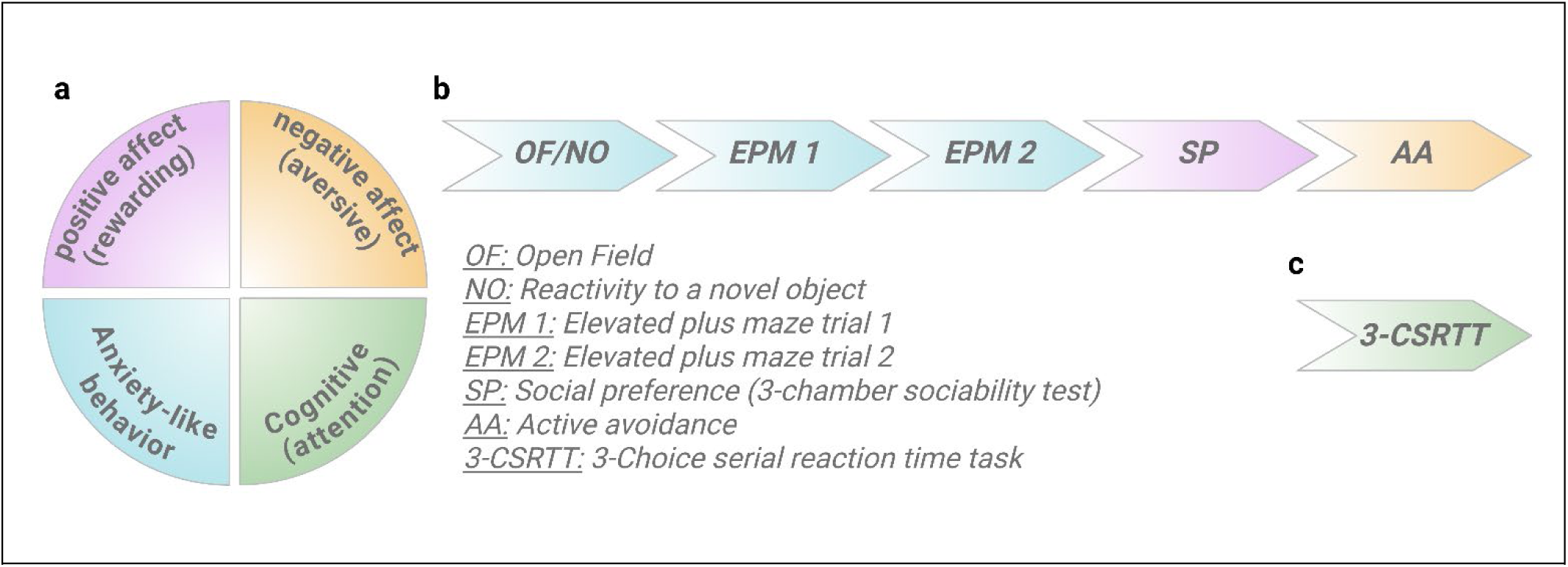
Schematic overview of behavioral investigation of dopaminergic *Hcrtr1* and *Hcrtr2* gene ablation. (**a**) Tests were selected across 4 axes: (1) positive affect (or rewarding experience) performed with the social preference test, (2) negative affect (or aversive experience) assessed via the electric shock active avoidance task, (3) anxiety-like behaviors assessed with the open field (OF), reactivity to novel object (NO), and elevated plus maze (EPM) tests and (4) cognitive behavior/attention assessment via the 3-choice serial reaction time task (3-CSRTT). (**b**) Temporal sequence of tests for anxiety-like, positive and negative affect behaviors. (**c**) The 3-CSRTT assessing attention performance was assessed with a different cohort of mice.

### Dopaminergic *Hcrtr1* ablation reduces explorative behavior and reactivity to novelty in an open field

We previously showed that the genetic inactivation of *Hcrtr2*, but not of *Hcrtr1*, nor the compound inactivation of both receptors in mouse DA neurons, caused a dramatic increase in time spent in the EEG theta-enriched, alert substate of wakefulness often referred to as ‘active wakefulness’, but in this case dissociated from locomotor activity^14,22^. In a reward-driven operant conditioning task based on visual discrimination, we found that the altered waking state of DA*^Ox2R-KO^* mice was associated with markedly increased learning rate and attentional skills compared to DA*^Ox2R-CT^* littermates. It however was also accompanied by deficits in impulse control, manifested by compulsive and impulsive-like behaviors^22^.

To functionally interrogate OX→DA neuromodulation in locomotor and explorative behaviors, we exposed mice to a 10-min open field test (‘OF’, **Fig. 5a, Left)**. While DA*^Ox1R-KO^* and DA*^Ox1R-CT^* mice traveled a similar total distance across the 45 x 45 cm arena (**Fig. 5c, Left**), DA*^Ox1R-KO^* mice spent more time in the wall zone and less time in the center zone respective to controls, suggesting increased thigmotaxis^42^ and anxiety-like behavior (**Fig. 5b-c**). Immediately following the 10-min OF test, a small object was placed in the middle of the center zone and mice were allowed a further 5 min of free exploration (novel object test ‘NO’, **Fig. 5a, Right**). Again, DA*^Ox1R-KO^* mice spent less time in the center zone of the OF, as well as in the center zone restricted to the object proximity (**Fig. 5e**). In contrast, DA*^Ox2R-KO^* and DA*^Ox2R-CT^* mice displayed no differences in total distance traveled, times spent in the wall or center zones (**Fig. S1a-b**), nor in exploration of the novel object (**Fig. S1c-d**). Altogether, our data reveal that DA*^Ox1R-KO^* mice show increased anxiety-like behavior during exploration of an OF, and reduced approach towards an object placed in the OF’s most anxiogenic zone.

**Figure 5.**
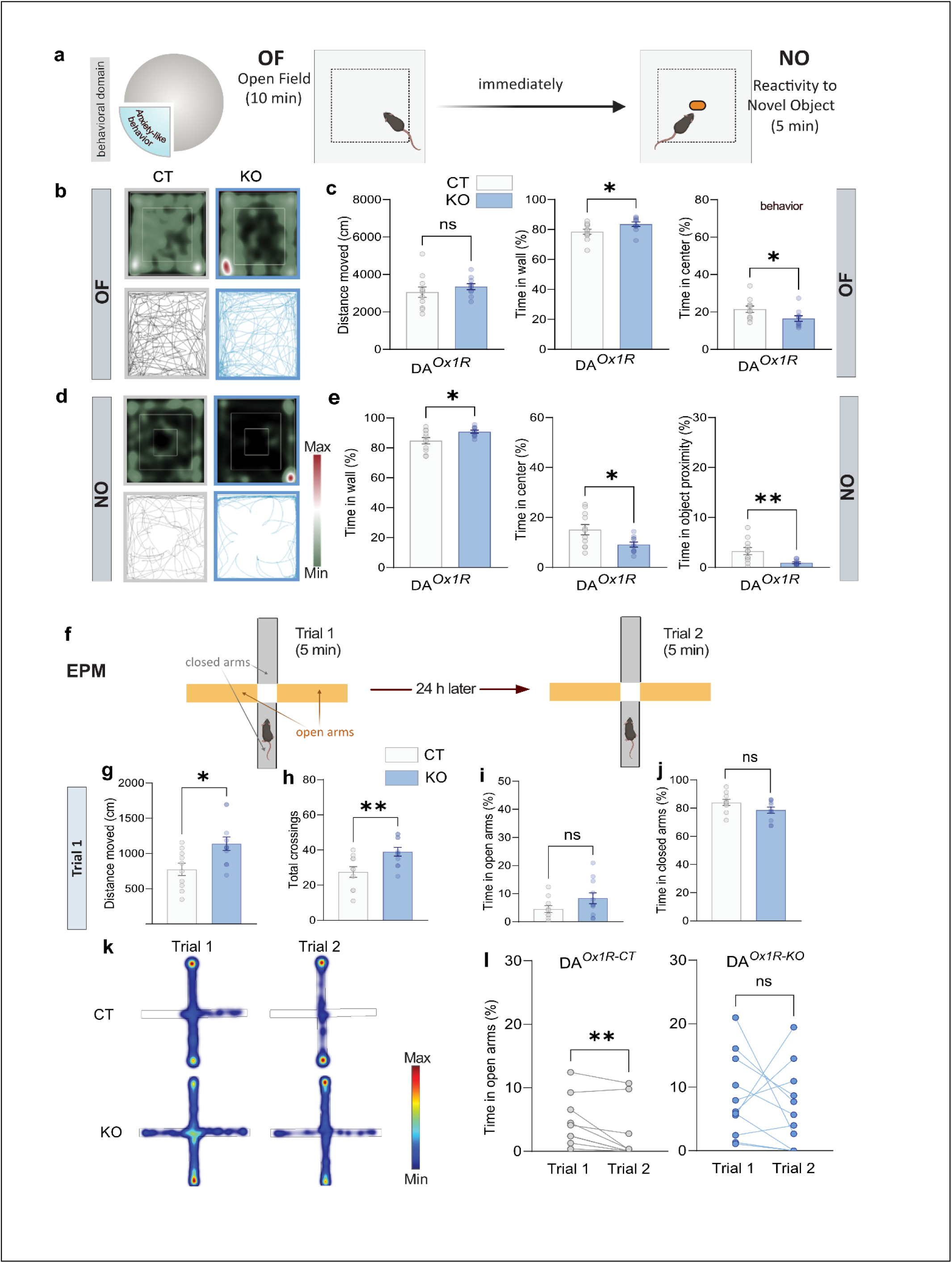
Dopaminergic *Hcrtr1*-ablated mice show reduced exploration and reactivity to novelty in an open field arena, but hyperactivity and atypical responding in an elevated plus maze. DA*^Ox1R-CT^* and DA*^Ox1R-KO^* mice were subjected to the open field (OF), reactivity to novel object (NO) and elevated plus maze (EPM) tests. (**a**) Schematic design of the OF and NO tests conducted in a 45 x 45 (cm) ground floor-sized arena. Mice were allowed to freely explore the open field for 10 min (OF). Immediately afterward, an object was placed in the arena center and mice allowed to freely explore the arena and the object for another 5 min (NO). The dashed-lined square delineates the center zone, defined as beyond 5.5 cm from the walls. (**b**) Heatmap (top) and trail map (bottom) of representative DA*^Ox1R-CT^* (left) and DA*^Ox1R-KO^* (right) mice during the 10 min OF exploration. Red and green indicate maximum and minimum occupancy, respectively. (**c**) (Left) Total distance traveled in the OF for DA*^Ox1R-CT^*and DA*^Ox1R-KO^* mice (*t*(19) = 0.911, *P* = 0.370, *t*-test); middle: time in the wall zone during OF (*t*(19) = 2.183, 0.042, *t*-test); (right) time in the center zone during OF (*t* (19) = 2.186, *P =* 0.041, *t*-test). **(d)** Heatmap (top) and trail map (bottom) of representative DA*^Ox1R-CT^* (left) and DA*^Ox1R-KO^* (right) mice during the 5 min NO test. (**e**) (Left) time in the wall zone during NO (*t*(19) = 2.513, *P =* 0.021, *t*-test); middle: time in the center zone during NO (*t*(19) = 2.523, *P =* 0.021, *t*-test); right: time in the center zone at 15 cm of the novel object (*t*(19) = 3.039, *P =* 0.007, *t*-test). *n* = 11 DA*^Ox1R-CT^*, 10 DA*^Ox1R-KO^*. (**f**) Schematic design of the EPM test. Mice were subjected to an EPM trial for 5 min, and 24 h later re-exposed to a 2^nd^ EPM trial for another 5 min. (**g**) Distance moved during the 1^st^ EPM trial revealed a higher total distance traveled by DA*^Ox1R-KO^* vs DA*^Ox1R-CT^* mice (*t* (19) = 2.777, *P =* 0.012, *t*-test), as well as (**h**) a higher total number of inter-zone crossings in KO vs CT (*t* (19) = 2.931, *P* = 0.009, *t*-test). (**i**) Time in the open arms (*t* (19) = 1.625, *P =* 0.121, *t*-test), and (**j**) time in the closed arms (*t* (19) = 1.790, *P* = 0.089, t-test) did not significantly differ in KO and CT. (**k**) Representative heatmaps depicting the location of a DA*^Ox1R-CT^* (top), and a DA*^Ox1R-KO^* mouse (bottom) during EPM trials 1 and 2. Heatmap occupancy color scale from red (Max) to dark blue (Min). (**l**) DA*^Ox1R-CT^* mice spent less time in open arms during the 2^nd^ EPM trial than during the 1^st^, as commonly observed in rodents (*W* = -51, *P* = 0.006, Wilcoxon test). In contrast, DA*^Ox1R-KO^* mice did not spend less time in open arms during the 2^nd^ EPM trial relative to the 1^st^ (*t* (10) = 0.652, *P =* 0.530, paired *t*-test). Bar graphs depict mean ± SEM. *n* = 10 DA*^Ox1R-CT^*, 11 DA*^Ox1R-KO^*.

### Dopaminergic *Hcrtr1*-ablated mice show hyperactivity and atypical behavior in the elevated plus maze

Because reduced exploration in the OF suggests heightened anxiety, we next evaluated DA*^Ox1R-KO^* mice in the elevated plus maze (EPM), a test commonly used to assess anxiety in rodents (**Fig. 5f**). While DA*^Ox1R-KO^*showed increases in total distance traveled through the maze (**Fig. 5g**), as well as increased total number of zone crossings relative to controls (**Fig. 5h**), no change in times spent in the open, or closed arms of the maze were observed during the 5-min test (**Fig. 5i-j**). Hence, loss of DA *Hcrtr1* induces context-dependent hyperactivity, but no reduced exploration of open arms. DA*^Ox1R-KO^* mice thus show misaligned anxiety-like behavioral responding in the EPM *vs* the OF.

Consistent with the OF test data, DA*^Ox2R-KO^* and DA*^Ox2R-CT^* mice did not differ in any EPM endpoint measures (**Fig. S1e-h**), indicating no alterations in locomotion, nor anxiety-like behavior.

The EPM test is considered to be a one-trial tolerance (i.e., non-repeatable) test, as WT mice exposed to a 2^nd^ EPM trial almost totally avoid the open arms^43,44,45^. However, because orexinergic modulation of DA targets was shown to play a critical role in risk assessment^46^ and executive control—particularly in impulsive and compulsive-like behaviors^22^—we asked whether a 2^nd^ EPM exposure would reveal behavioral changes indicative of alterations in risk assessment, impulse control, or phobic-like avoidance of the open arms^47,48^. Thus, 24 h after the 1^st^ EPM trial, we exposed mice to a 2^nd^ trial. As expected from prior reports, the total distance traveled was reduced between the 1^st^ and 2^nd^ trials in all 4 genotype groups (DA*^Ox1R-CT^*, DA*^Ox1R-KO^*, DA*^Ox2R-CT^*, and DA*^Ox2R-KO^*; **Fig. S2**). Interestingly, however, whereas DA*^Ox1R-CT^* mice show, as expected, decreased time spent in the open arms during the 2^nd^ trial compared to the 1^st^, DA*^Ox1R-KO^* mice did not show reduced open arms occupancy between trials 1 and 2 (**Fig. 5k-l**). In contrast, both DA*^Ox2R-KO^* and DA*^Ox2R-CT^* showed the expected pattern of reduced time in open arms during the 2^nd^ trial relative to the 1^st^ (**Fig. S1i-l**).

In sum, our findings support a specific role of DA OX_1_R signaling, but not OX_2_R, for context-dependent hyperactivity and anxiety-like behavior. Moreover, because DA*^Ox1R-KO^*do not show reduced open arm occupancy in EPM trial 2 vs trial 1, our data suggest that DA cell OX_1_R signaling may have a function in behavioral adaptation, acquisition of phobic-like responses and/or impulse control.

### Dopaminergic *Hcrtr1*-ablated mice exhibit increased impulsive and compulsive responding in a reward-driven operant task

To follow up on our EPM findings and further delineate their behavioral profile, DA*^Ox1R-KO^* mice were next exposed to an operant conditioning paradigm. Specifically, we used the 3-Choice Serial Reaction Time Task (3-CSRTT), a learning task equivalent to the Continuous Attention Performance Task in humans, measuring sweet reward-driven instrumental learning, attentional skills, compulsive and impulsive-like traits^49^ (**Fig. 6a**). As mentioned above, in a similar task, DA*^Ox2R-KO^* mice exhibited markedly enhanced learning rate, and, once performance was stabilized and test contingencies were made more difficult, profoundly improved response accuracy, indicating higher attentional skills. Improved performance was however accompanied by maladaptive patterns of reward-seeking, with compulsive and impulsive-like responses^22^.

**Figure 6.**
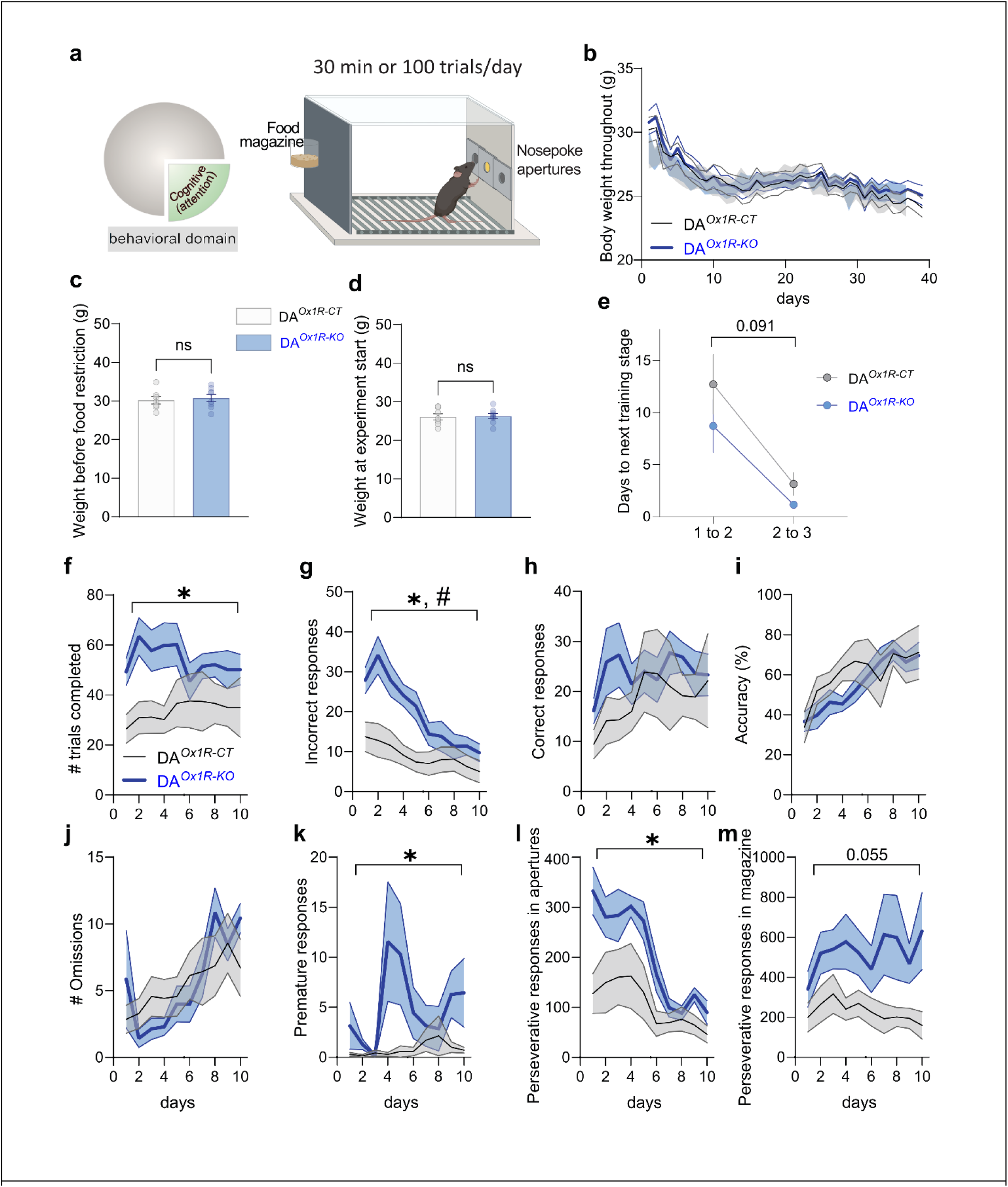
Dopaminergic *Hcrtr1*-ablated mice exhibit hyperactivity, impulsive- and compulsive-like behaviors in reward-driven operant conditioning. Performance of DA*^Ox1R-KO^* (KO) and DA*^Ox1R-CT^*(CT) mice in the 3-choice serial reaction time task (3-CSRTT). (**a**) Schematic of the 3-CSRTT operant chamber. Three nosepoking apertures were located on the wall opposite the food magazine dispensing chocolate-flavored sucrose pellet rewards. (**b**) Timeline of body weight throughout the 3-CSRTT procedure showed no difference between KO and CT mice (time: *F*_38,494_ = 19.881, *P <* 0.001; genotype: *F*_1,13_ = 0.061, *P* = 0.808; time × genotype: *F*_38,494_ = 0.363, *P =* 1.000, repeated measures 2-way ANOVA). (**c**) The body weight of the mice before food restriction (*t* (13) = 0.417, *P =* 0.683, *t*-test), and (**d**) the body weight at the start of the 3-CSRTT procedure (*t* (13) = 0.200, *P =* 0.844, *t*-test) did not differ between KO and CT mice, *n* = 7 DA*^Ox1R-CT^*, 8 DA*^Ox1R-KO^*. (**e**) DA*^Ox1R-KO^* mice showed a tendency to learn faster during the initial training days (training stages 1-3) relative to controls, as shown by comparing number of days to reach the next training stage in DA*^Ox1R-KO^* and DA*^Ox1R-CT^*mice (time: *F*_1,12_ = 15.517, *P =* 0.002, genotype: *F*_1,12_ = 1.827, *P =* 0.201, time × genotype: *F*_1,12_ = 3.379, *P =* 0.091, repeated-measures 2-way ANOVA). (**f**) DA*^Ox1R-KO^* completed a higher number of total trials compared to DA*^Ox1R-CT^* littermates [time: *F*_2.625,31.497_ = 0.653, *P=* 0.567; genotype: *F*_1,12_ = 5.235, *P =* 0.041; time × genotype: *F*_2.625,31.497_ = 1.183, *P =* 0.328, repeated-measures 2-way ANOVA], as well as (**g**) a higher number of incorrect responses during the initial 10 training days (time: *F* (3.551, 42.611) = 14.670, *P <* 0.001; genotype: *F*_1,12_ = 8.389, *P =* 0.013, time × genotype: *F* (3.551, 42.611) = 4.413, *P =* 0.006, repeated-measures 2-way ANOVA). DA*^Ox1R-KO^* mice, however, did not differ from DA*^Ox1R-CT^* mice in (**h**) number of correct responses (time: *F*_1.977,23.719_ = 1.614, *P =* 0.220, genotype: *F*_1,12_ = 0.863, *P =* 0.371, time × genotype: *F*_1.977,23.719_ = 0.762, *P =* 0.476, repeated-measures 2-way ANOVA), or (**i**) response accuracy (time: *F*_2.372,28.461_ = 8.731, *P <* 0.001, genotype: *F*_1,12_ = 0.320, *P =* 0.582, time × genotype: *F*_2.372,28.461_ = 1.258, *P =* 0.304, repeated measures 2-way ANOVA), or (**j**) number of omissions (time: *F*_2.630,31.563_ = 5.033, *P =* 0.008; genotype: *F*_1,12_ = 0.009, *P =* 0.928; time × genotype: *F*_2.630,31.563_ = 1.380, *P =* 0.268, repeated-measures 2-way ANOVA) during the first 10 training days. (**k**) DA*^Ox1R-KO^* mice exhibited a higher number of premature responses compared to DA*^Ox1R-CT^* mice during the 1^st^ 10 training days (time: *F*_2.175,26.100_ = 1.573, *P =* 0.226, genotype: *F*_1,12_ = 5.775, *P =* 0.033, time × genotype: *F*_2.175,26.100_ = 1.767, *P =* 0.189, repeated-measures 2-way ANOVA), (**l**) a higher number of repetitive nosepoking responses in any of the 3 apertures (time: *F*_3.456,41.468_ = 13.310, *P <* 0.001, genotype: *F*_1,12_ = 6.294, *P =* 0.027, time × genotype: *F* _3.456,41.468_ = 2.478, *P =* 0.067, repeated-measures 2-way ANOVA), and (**m**) a trend towards a higher number of repetitive nosepoking responses in the food magazine (time: *F*_2.753,33.031_ = 1.451, *P =* 0.247, genotype: *F*_1,12_ = 4.501, *P =* 0.055, time × genotype: *F*_2.753,33.031_ = 1.677, *P =* 0.194, repeated-measures 2-way ANOVA). Bar graphs and timelines depict mean ± SEM. *n* = 7 DA*^Ox1R-CT^*, 7 DA*^Ox1R-KO^*. * denotes a significant genotype effect, and # a significant time × genotype effect, in repeated-measures 2-way ANOVA.

As the 3-CSRTT procedure comprises mild food restriction, body weight was monitored throughout the procedure. No weight differences between DA*^Ox1R-KO^* and DA*^Ox1R-CT^* mice were observed, either before food restriction (**Fig. 6c**), at experimental start (**Fig. 6d**) or throughout the learning process (**Fig. 6b**). DA*^Ox1R-KO^* and DA*^Ox1R-CT^* mice displayed similar learning rates from one stage to the next (**Fig. 6e**). However, during the first 10 training days, DA*^Ox1R-KO^* mice completed a higher number of total trials (**Fig. 6f**), a higher number of premature responses (**Fig. 6k**), and a higher number of incorrect responses (**Fig. 6g**). No differences were observed in correct responses, response accuracy or response omissions relative to controls (**Fig. 6h-j**). In addition, compared to controls, DA*^Ox1R-KO^* mice showed an increased number of repetitive responses in the nosepoking apertures (**Fig. 6l**), and a trend toward higher repetitive nosepoking in the food magazine (**Fig. 6m**). Repetitive nosepoking responses in the apertures, or in the food magazine, are considered indices of compulsive behavior^49^.

The data described above assess performance in early training when individual mice may be in different training stages depending on performance. However, when the mice were assessed at the same training stages after adequate learning, no differences were observed between groups for either easy (Stage 3; **Fig. S3a-e**), or more challenging (Stage 5; **Fig. S3f-j**) task contingencies. Interestingly, these data indicate that differences in performance between DA*^Ox1R-KO^* and DA*^Ox1R-CT^*are evident in early phase of 3-CSRTT training but are occluded as training progresses.

Considering the present data of DA*^Ox1R-KO^* mice and prior DA*^Ox2R-KO^* mice’ data^22^, either DA OX_1_R or DA OX_2_R loss leads to compulsive behavioral patterns, suggesting that DA OXR signaling is protective against such traits. Although both mutants show compulsive-like behavior, this trait manifests in the context of pronounced cognitive improvement, i.e., higher learning and response accuracy, solely in DA*^Ox2R-KO^* mice.

### Dopaminergic *Hcrtr1*-ablated mice show normal aversion-driven instrumental learning

After evaluating reward-driven learning, we next assessed associative learning based on negative affect using the active avoidance task (**Fig. 7a**). This 2-day task assays the mouse’s ability to learn to avoid an electric shock preceded by a 10-s-lasting tone by transiting to an adjacent chamber. DA*^Ox1R-KO^* and controls displayed similar learning performance, i.e., similar numbers of shock avoidances (i.e., transiting to the safe chamber during the shock-predicting tone, considered as index of successful learning, **Fig. 7d**), escapes (i.e., transitions to the other chamber after the shock had commenced, **Fig. 7f**), and premature responses (transitions to the chamber before the tone had started, **Fig. 7b**). Whereas latencies for premature responses on day 1 and 2 did not differ between KO and CT (**Fig. 7c**), DA*^Ox1R-KO^* mice showed a shorter latency to avoid the shock on day 1 (**Fig. 7e**). No differences in escape latencies were observed between KO and CT on either of the 2 days (**Fig. 7g**). Finally, all mice from the DA*^Ox1R-CT^* group learnt the task while 1 out of the 11 DA*^Ox1R-KO^* mice tested was classified as a non-learner (**Fig. 7h-j**; see Methods for classification criteria).

**Figure 7.**
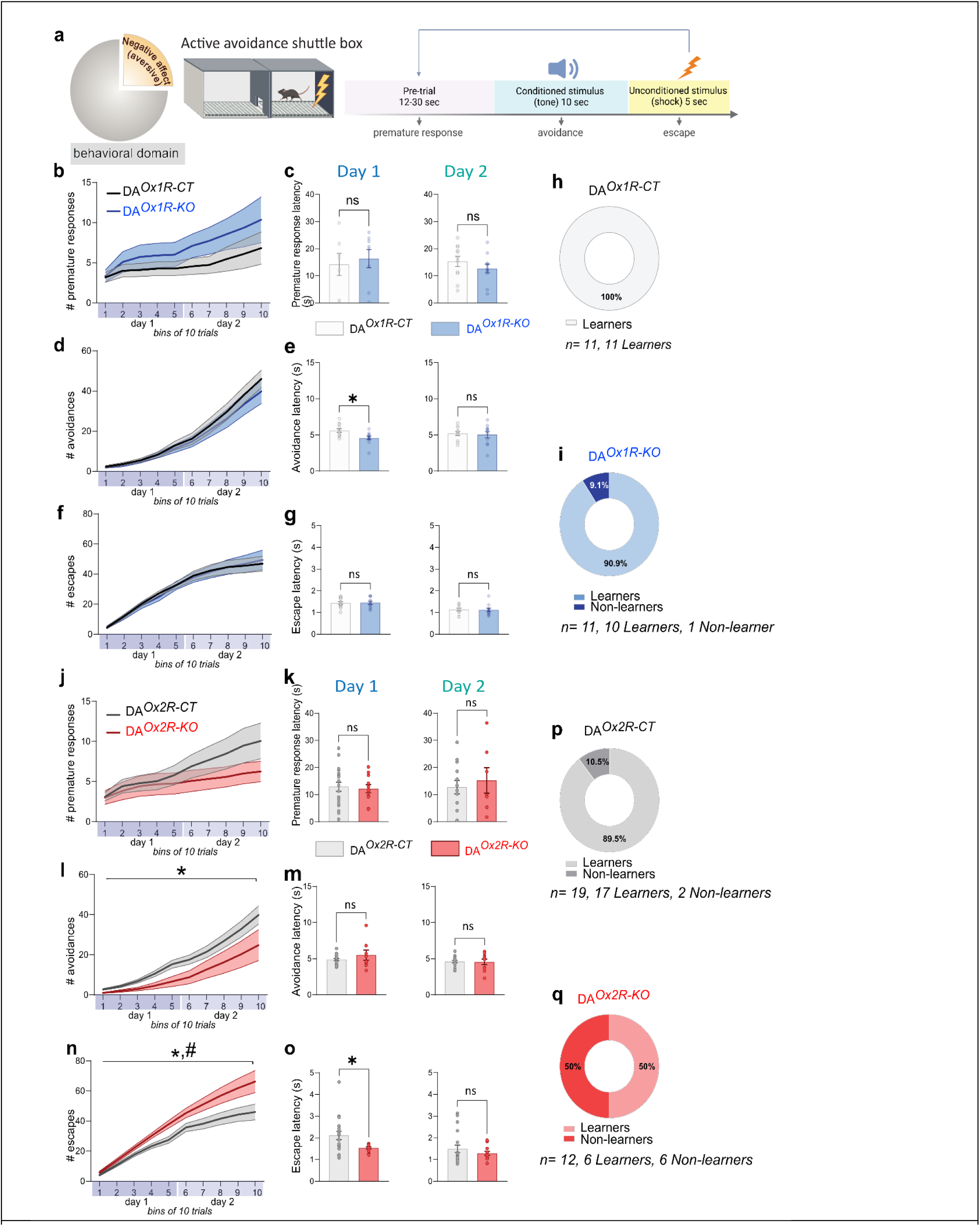
Dopaminergic *Hcrtr2*-ablated mice show impaired aversion-driven instrumental learning. Performance of DA*^Ox1R-KO^* vs DA*^Ox1R-CT^*and DA*^Ox2R-KO^* vs DA*^Ox2R-CT^* mice in an active avoidance task. (**a**) Schematic of the active avoidance shuttle box and experimental design. An inter-chamber run during the pre-trial is a premature response, an inter-chamber run during the tone is an avoidance response (indicative of successful learning), an inter-chamber run during the shock is an escape response. (**b**) The number of premature responses did not differ between DA*^Ox1R-CT^* and DA*^Ox1R-KO^* mice across the 2 days of testing (time: *F*_1.233,24.668_ = 7.511, *P <* 0.001, genotype: *F*_1,20_ = 1.374, *P =* 0.255, time × genotype: *F*_1.233,24.668_ = 0.878, *P =* 0.380, repeated-measures 2-way ANOVA). (**c**) Similar premature response latencies in DA*^Ox1R-CT^* and DA*^Ox1R-KO^* mice (Day 1, left: *t* (13) = 0.400, *P =* 0.695; Day 2, right: *t* (20) = 1.082, *P =* 0.292; t-tests). (**d**) The number of avoidance responses did not differ between DA*^Ox1R-CT^*and DA*^Ox1R-KO^* mice across the 2 days of testing (time: *F*_1.126, 22.519_ = 97.899, *P <* 0.001; genotype: *F*_1,20_ = 0.369, *P =* 0.550; time × genotype: *F*_1.126, 22.519_ = 0.502, *P =* 0.507, repeated-measures 2-way ANOVA). (**e**) DA*^Ox1R-KO^* mice showed shorter avoidance response latencies than DA*^Ox1R-CT^* in the 1^st^ day of testing, indicating a faster avoiding response (Day 1, left: *t* (20) = 2.796, *P =* 0.011, Day 2, right: *t* (20) = 0.290, *P =* 0.775; t-tests). (**f**) Number of escape responses did not differ between DA*^Ox1R-CT^* and DA*^Ox1R-KO^* mice across the 2 days of testing (time: *F*_1.156,23.113_ = 102.588, *P <* 0.001; genotype: *F*_1,20_ = 0.003, *P =* 0.960; time × genotype: *F*_1.156,23.113_ = 0.200, *P =* 0.695; repeated-measures 2-way ANOVA). (**g**) Similar escape response latencies in DA*^Ox1R-CT^* and DA*^Ox1R-KO^* mice (Day 1, left: *t* (20) = 0.048, *P =* 0.962, and Day 2, right: *U* = 51.50, *P =* 0.573 t-test and Mann-Whitney test, respectively). (**h**) and (**i**) Pie charts of learners *vs* non-learners in DA*^Ox1R-CT^*and DA*^Ox1R-KO^* show that both genotypes learn efficiently. *n* = 11 DA*^Ox1R-CT^*, 11 DA*^Ox1R-KO^*. (**j**) The number of premature responses did not differ between DA*^Ox2R-CT^* and DA*^Ox2R-KO^* across the 2 days of testing (time: *F*_1.303,37.779_ = 12.325, *P <* 0.001, genotype: *F*_1,29_ = 0.744, *P =* 0.396, time × genotype: *F*_1.303,37.779_ = 2.178, *P =* 0.143, repeated-measures 2-way ANOVA) (**k**) Similar premature response latencies in DA*^Ox2R-CT^* and DA*^Ox2R-KO^* mice (Day 1, left: *t* (28) = 0.290, *P =* 0.774; Day 2, right: *t* (18) = 0.528, *P =* 0.604; *t*-tests). (**l**) DA*^Ox2R-KO^* mice show a decreased number of successful avoidance responses compared to DA*^Ox2R-CT^* mice across the 2 days of testing (time: *F*_1.146,33.242_ = 46.938, *P <* 0.001, genotype: *F*_1,29_ = 4.507, *P =* 0.042, time × genotype: *F*_1.146,33.242_ = 2.084, *P =* 0.156, repeated-measures 2-way ANOVA). (**m**) Similar avoidance response latencies in DA*^Ox2R-CT^* and DA*^Ox2R-KO^* mice (Day 1, left: *t* (24) = 1.185, *P =* 0.248; Day 2, right: *t* (25) = 0.111, *P =* 0.912; *t*-tests). (**n**) DA*^Ox2R-KO^* mice show an increased number of escapes (inter-chamber run after shock exposure) compared to DA*^Ox2R-CT^* mice, with both a genotype and a time x genotype significant effect (time: *F*_1.286, 37.284_ = 126.005, *P <* 0.001; genotype: *F*_1,29_ = 6.354, *P =* 0.017; time × genotype: *F*_1.286, 37.284_ = 3.833, *P =* 0.048, repeated-measures 2-way ANOVA). (**o**) DA*^Ox2R-KO^* mice nevertheless show shorter escape response latencies than DA*^Ox2R-CT^*mice in the 1^st^ day of testing, suggesting that they are sensitive to the electric shock (Day 1, left: *U* = 62, *P =* 0.035; Day 2, right: *U* = 113, *P =* 0.984; Mann-Whitney tests). (**p**) and (**q**) Pie charts of learners *vs* non-learners in DA*^Ox2R-CT^* and DA*^Ox2R-KO^* mice highlight the markedly reduced learning efficiency observed among DA*^Ox2R-KO^*mice. Bar graphs and timelines depict mean ± SEM. *n* = 19 DA*^Ox2R-CT^*, 12 DA*^Ox2R-KO^*.

In sum, our findings suggest that while loss of OX_1_R-mediated modulation of DA cells compromises impulse control in a positive affect-driven operant conditioning task, leading to impulsive and compulsive responses, negative affect-driven learning in an active avoidance task appears unaffected, with the exception that mice exhibit faster reaction times in shock-avoidance responding than do controls. This feature may be another indicator of an impulsive component, in line with premature responding in the reward-driven task, or reflect altered risk assessment, in line with absence of diminished open arm exploration in the 2^nd^ EPM trial.

### Dopaminergic *Hcrtr2*-ablated mice show impaired aversion-based instrumental learning

Previously, we showed that DA*^Ox2R-KO^* mice exhibited faster learning and markedly improved attentional skills in a reward-driven visual-discrimination-based operant conditioning task^22^. Here, when tested in an avoidance-driven instrumental learning task, we found that these mice exhibited a severely compromised performance. Specifically, in the active avoidance test, DA*^Ox2R-KO^*mice showed a lower number of successful avoidances, i.e., transitions to the adjacent chamber during the conditioned stimulus (tone) and before electric shock start (**Fig. 7l**), and incurred more electric shocks, as they showed a higher number of escapes, i.e., inter-chamber runs after completion of the 10-s tone conditioned stimulus and the electric shock start (**Fig. 7n**). No differences in premature responding were observed (**Fig. 7j**).

Because task learning is here based on pain sensitivity, impaired active avoidance could be secondary to a sensory deficit (Razavi 2017). Our data suggest this interpretation however to be unlikely since, compared to controls, DA*^Ox2R-KO^* mice exhibited shorter escape latency in day 1 of learning (**Fig. 7o**), and a higher number of escapes throughout the two test days (**Fig. 7n**) suggesting that DA*^Ox2R-KO^*mice do perceive the electric shock as an aversive stimulus, but do not learn to integrate the test contingencies as efficiently as control littermates. When classifying the mice into learners and non-learners, only 2/19 control mice classified as non-learners (**Fig. 7p**), whilst 6/12 DA*^Ox2R-KO^* mice classified as non-learners (**Fig. 7q**). Hence, the constitutively hyperaroused EEG of DA*^Ox2R-KO^* mice appears to put them at an advantage in attention for positively-valued stimuli^22^, but not for negatively-valued stimuli.

### Dopaminergic *Hcrtr2*-ablated mice show reduced sociability

Given the important role of DA^50–52^ and OX^53,54^ neurons in social behavior, we asked whether loss of *Hcrtr1* or *Hcrtr2* from DA neurons alters the social interaction domain and approach behavior. In the three-chamber sociability test^55^, after 10 min of free exploration and habituation to the arena, mice can explore an empty enclosure, or an enclosure containing a non-familiar juvenile mouse (**Fig. 8a**). We found that DA*^Ox1R-KO^* mice manifest increased locomotion compared to controls during the habituation phase (when no enclosures were present). This manifested with an increased total distance traveled (**Fig. 8b**), and increased number of inter-chamber-crossings of DA*^Ox1R-KO^* vs DA*^Ox1R-CT^* mice (**Fig. 8c**). During the test, however both DA*^Ox1R-KO^* and DA*^Ox1R-CT^* mice showed intact sociability, with a larger % of time spent in proximity of the juvenile mouse than the empty enclosure (**Fig. 8e-f**), and a longer average exploration bout duration around the social than the inanimate stimulus (**Fig. 8g-h**), with no difference in total distance traveled during this part of the test (**Fig. S4**).

**Figure 8.**
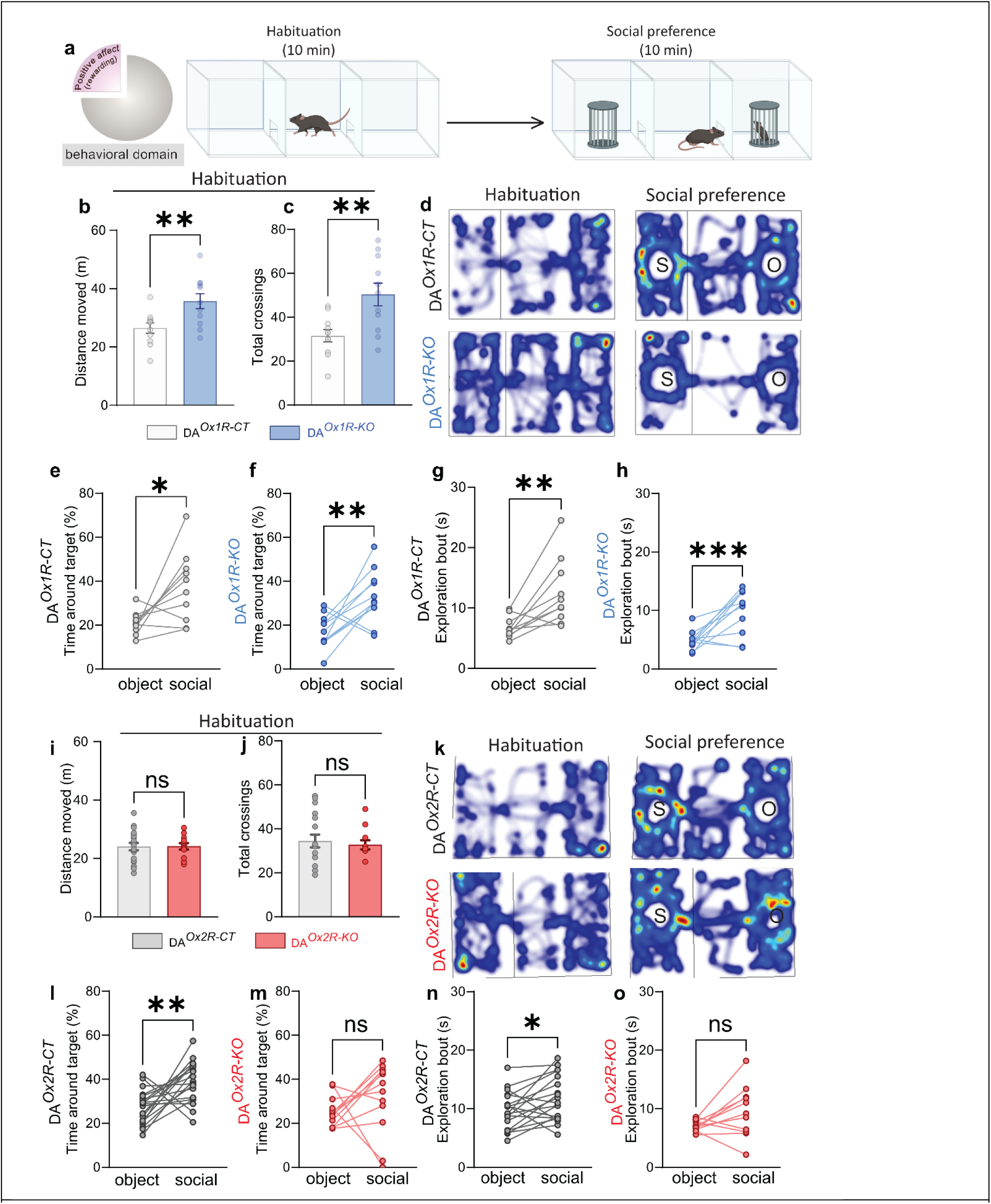
Dopaminergic *Hcrtr2*-ablated mice show reduced sociability. The social preference of DA*^Ox1R-KO^* and DA*^Ox2R-KO^* mice was probed against their respective genetic controls using the 3-chamber test. (**a**) Schematic of the experimental design consisting of two phases (i) a 10 min-habituation phase, during which the mouse is allowed to freely explore the arena, and (ii) a subsequent 10 min-social preference phase, during which two grided enclosures are located in the two opposite chambers, one empty and one containing a same-sex juvenile non-familiar mouse, and the experimental mouse is allowed to freely explore the 3 chambers and interact with the juvenile using olfactory, visual, auditory, and tactile cues. During the habituation phase, DA*^Ox1R-KO^* mice were found to (**b**) travel across a longer total distance (*t*_20_ = 2.972, *P =* 0.007, *t*-test, *n* = 11 DA*^Ox1R-CT^*, 11 DA*^Ox1R-KO^*), and (**c**) perform a higher number of inter-zone crossings relative to DA*^Ox1R-CT^* mice (*t*_20_ = 3.224, *P =* 0.004, *t*-test, *n* = 11 DA*^Ox1R-CT^*, 11 DA*^Ox1R-KO^*). (**d**) Representative heatmaps of the location of a DA*^Ox1R-CT^* and a DA*^Ox1R-KO^* mouse during the habituation (left) and the social preference (right) phases. “S” denotes the social stimulus-containing enclosure, “O” denotes the object (empty) enclosure. Both DA*^Ox1R-CT^*(**e**) and DA*^Ox1R-KO^*(**f**) mice spent a higher % of total time within proximity of the social than inanimate target (DA*^Ox1R-CT^*: *t* (9) = 2.840, *P =* 0.019; DA*^Ox1R-KO^*, *t* (10) = 3.230, *P =* 0.009; paired *t*-test). Both DA*^Ox1R-CT^* (**g**) and DA*^Ox1R-KO^* (**h**) mice displayed longer exploration bouts within proximity of the social than inanimate target (DA*^Ox1R-CT^*: *t*_9_ = 3.564, *P =* 0.006; for DA*^Ox1R-KO^*: *t* (10) = 3.987, *P =* 0.003; paired *t*-test). *n* = 10 DA*^Ox1R-CT^*, 11 DA^Ox1R-KO^). In contrast to DA*^Ox1R-KO^* mice, during the habituation phase, DA*^Ox2R-KO^* mice did not exhibit any differences in total distance traveled (**i**), nor in total number of inter-chamber-crossings (**j**), compared to DA*^Ox2R-CT^* control mice (total distance moved: DA*^Ox2R-CT^* vs DA*^Ox2R-KO^*, *t* (29) = 0.065, *P =* 0.949, t-test, *n* = 19 DA*^Ox2R-CT^*, 12 DA*^Ox2R-KO^*; total inter-chamber crossings: DA*^Ox2R-CT^* vs DA*^Ox2R-KO^*, *U* = 110.5, *P =* 0.897, Mann-Whitney test, *n* = 19 DA*^Ox2R-CT^*, 12 DA*^Ox2R-KO^*). (**k**) Representative heatmaps of a DA*^Ox2R-CT^* and a DA*^Ox2R-KO^* mouse during the habituation phase (left) and during social preference (right). Analysis of social preference in DA*^Ox2R-CT^* and DA*^Ox2R-KO^*mice revealed that while control mice show normal social preference (**l**, time around social and empty enclosures for DA*^Ox2R-CT^*, *t* (18) = 3.386, *P =* 0.003, paired *t*-test), and longer exploration bouts of the social than inanimate target (**n**, average exploration bout duration DA*^Ox2R-CT^* mice: *t* (18) = 2.393, *P =* 0.028, paired *t*-test), DA*^Ox2R-KO^* mice neither exhibited social preference in % time spent around the juvenile mouse (**m**, paired comparison of time around the social vs empty enclosures for DA*^Ox2R-KO^*, *W* = 30.00, *P =* 0.266, Wilcoxon test), nor manifested longer mean exploration bout duration for the social vs inanimate target (**o**, paired comparison of average exploration bout duration for DA*^Ox2R-KO^*, *t* (10) = 1.927, *P =* 0.083). Bar graphs show individual values and mean ± SEM. *n* = 19 DA*^Ox2R-CT^*, 12 DA*^Ox2R-KO^*.

**Figure 9.**
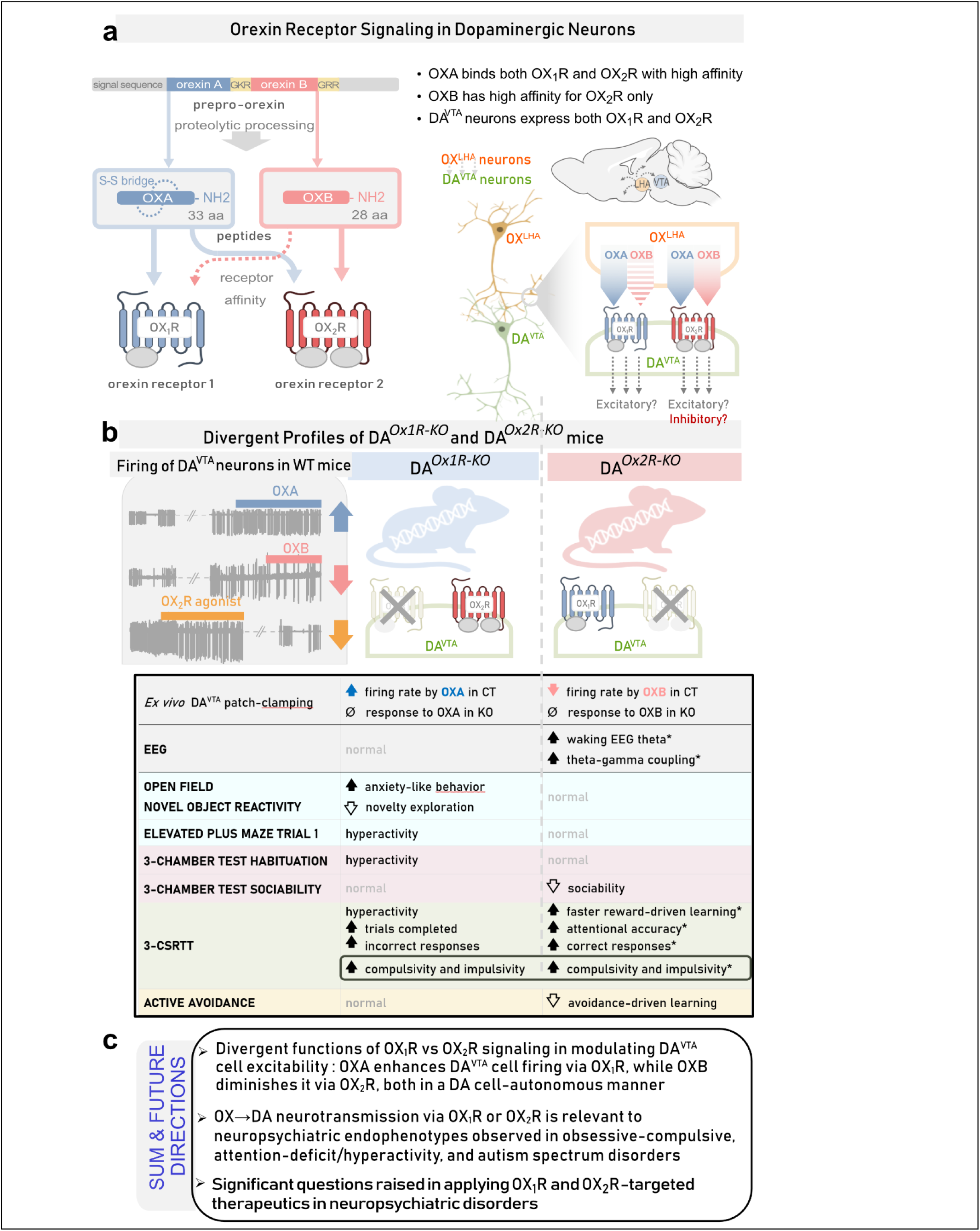
Summary depiction of divergent neuromodulation of dopaminergic neurons by hypocretin/orexin receptors-1 and -2 as reflected at electrophysiological and behavioral levels. (**a**) OX^LHA^→DA^VTA^ neurotransmission. Proteolytic cleavage of a common prepro-orexin precursor gives rise to Orexin A (OXA) and Orexin B (OXB) peptides. OX cell bodies in the lateral hypothalamic area (LHA) project to the ventral tegmental area (VTA) where they bind two G-protein-coupled receptors, OX_1_R and OX_2_R, both expressed in DA^VTA^ neurons^22^. OXA binds OX_1_R and OX_2_R with high affinity. OXB only shows high affinity for OX_2_R. (**b**) DA-specific genetically engineered knockout (KO) mouse models. DA*^Ox1R-KO^* mice lack the *Hcrtr1* gene in DA cells, making them unresponsive to OX_1_R input. DA*^Ox2R-KO^* mice lack the *Hcrtr2* gene in DA cells, eliminating OX_2_R responses. Control groups (DA*^Ox1R-CT^* and DA*^Ox2R-CT^*) are littermates with intact receptor expression. Electrophysiological recordings show that OXA enhances firing of DA^VTA^ neurons from DA*^Ox1R-CT^* but not DA*^Ox1R-KO^* mice, confirming OX_1_R-dependent excitation by OXA. In contrast, OXB decreases firing of DA^VTA^ cells from DA*^Ox2R-CT^* mice, but not DA*^Ox2R-KO^* mice, indicating OX_2_R-dependent inhibition by OXB. Behaviorally DA*^Ox1R-KO^* mice exhibit increased anxiety-like behavior, hyperactivity in the elevated-plus-maze (EPM) and the 3-chamber apparatus during the habituation phase, and hyperresponding in the 3-CSRTT. DA*^Ox2R-KO^* mice display markedly increased EEG waking theta and enhanced theta-gamma coupling, improved learning rate and attentional skills in reward-driven 3-CSRTT. They exhibit severely compromised aversion-driven instrumental learning and reduced sociability. Both DA*^Ox1R-KO^* and DA*^Ox2R-KO^* mice exhibit enhanced impulsivity and compulsive-like behaviors. Summary, Open Questions & Clinical Relevance. Electrophysiology shows that OXA enhances DA^VTA^ neuronal excitability via OX_1_R, while OXB diminishes it via OX_2_R, both in a DA cell-autonomous manner. These opposing cellular actions lead to distinct behavioral outcomes. The study positions OX->DA neurotransmission via OX_1_R or OX_2_R as relevant to endophenotypes observed in obsessive-compulsive, attention-deficit/hyperactivity, and autism spectrum disorders, raising important questions for pharmacological therapy, as multiple selective or dual-OX_1_R and OX_2_R-targeted therapeutics are being considered. Asterisks denote data from Bandarabadi *et al*, 2023^22^.

In contrast to DA*^Ox1R-KO^* mice, DA*^Ox2R-KO^* mice manifested no hyperlocomotion, i.e., no differences in total distance traveled (**Fig. 8i**), nor in number of inter-chamber crossings (**Fig. 8j**) during the habituation phase relative to DA*^Ox2R-CT^* controls. Remarkably, however, while DA*^Ox2R-CT^* mice showed a clear preference for the juvenile mouse vs the empty enclosure, DA*^Ox2R-KO^* mice did not (**Fig. 8k-m**). Additionally, DA*^Ox2R-CT^* mice showed longer mean exploration bout durations for the juvenile mouse rather than for the object (**Fig. 8n**), but DA*^Ox1R-KO^* did not (**Fig. 8o**). Total distance traveled did not differ (**Fig. S4**). In sum, our data newly reveal a critical role of OX_2_R-mediated modulation of DA neuronal activity for expression of social preference in mice.

## DISCUSSION

OX and DA are both key neural hubs involved in processing past and present salient information, essential for the adaptive regulation of arousal, motivation, and stress responses. Their interactions remain however poorly defined due to difficulties in simultaneously interrogating their functions *in vivo*. OX^LH^ cells send dense projections to DA^VTA^ neurons and a body of data supports OX^LH^→DA^VTA^ connectivity as critical for behaviors associated with reward- and aversion-paired stimuli relevant to motivational states and related-neuropsychiatric disorders^7,26,27,28–31^. In this pathway, little is known about cell-specific OX_1_R *vs* OX_2_R contribution. Our study aims to fill some of these gaps.

To probe the functional roles of the two receptors in OX→DA connectivity, we generated mice whose DA neurons are unable to respond to OXs by selective genetic disruption of *Hcrtr1*, *Hcrtr2*, or both, in DA cells^22^. Analysis of the mice’ EEG activity and sleep/wake vigilance state architecture revealed that DA-specific *Hcrtr2* loss caused a dramatic increase in time spent in a constitutively high-alertness type of wakefulness with enhanced EEG theta-gamma coupling. This was paired with markedly improved performance in a continuous attention operant task, albeit also with compulsive and impulsive behaviors^22^. DA-specific *Hcrtr1* loss did not cause similar changes in EEG-dependent parameters, while coupled loss of *Hcrtr1* and *Hcrtr2* in DA neurons, tended to cause opposite changes, with a wakefulness displaying decreased theta power^22^.

Here we extend our study of the contribution of OX→DA pathways to a wider range of behavioral domains. Additionally, based on evidence pointing to modulation of DA^VTA^ neurons as central component of OX/DA interactions, we interrogate the underlying DA^VTA^ cell electrophysiological substrates. We discovered that inactivating OX_1_R or OX_2_R in DA cells both profoundly affected DA^VTA^ cell excitability and mouse behavioral profile, but in strikingly different ways, thus uncovering for the first time divergent orexinergic modulation of dopaminergic function by OX_1_R *vs* OX_2_R.

### DA*^Ox1R-KO^* mouse behavior: anxiety and hypermotricity

DA*^Ox1R-KO^* mice displayed context-specific motor hyperactivity, accompanied by marked impulsivity and repetitive behaviors. This was manifested by increased total distance traveled and inter-zone crossings in both the EPM and the 3-chamber arena, and by increased total trials completed, premature, incorrect, and perseverative responses during the 3-CSRTT.

Explorative behavior was context-dependently altered. While DA*^Ox1R-KO^*mice engaged in increased exploration and locomotion in confined arenas, such as the EPM or 3-chamber arena, exploration and novel object approach in large open areas, such as the OF (45 x 45 cm), were reduced, suggesting anxiety-like behavior. While the EPM is one of the most commonly used tests to study anxiety-like behavior in rodents^56^, the finding of anxiety-like responding detected in the OF was not paralleled in the EPM. OF exploration implicates locomotor, motivational and emotional aspects^30^ that differ from EPM exploration. Dissociation between OF and EPM mouse responding was reported by others^57^. Context-dependent differences in anxiety-like read-outs may have multiple causes. First, the OF was the 1^st^ paradigm our mice were subjected to in our behavioral pipeline (**Fig. 4**). Despite the preliminary handling procedure adopted to habituate mice to the experimenter, anxiety-like behavior may diminish as mice gradually acclimate to experimental testing. Second, the two tests measure different aspects of anxiety-like behavior^58^. Third, variables such as task duration (10 min in OF, 5 min in EPM) or other differences in spatiotemporal configuration of the tests may cause divergent test readouts. The EPM one-trial tolerance effect^43^ was shown to involve NMDAR activation and D_1_R suppression in the hippocampus^45^. Its occlusion in DA*^Ox1R-KO^*mice suggests that these pathways may lie downstream of DA OX_1_R signaling.

Our finding of reduced OF exploration in DA*^Ox1R-KO^* mice misaligns with reports by Xiao *et al*^59^ describing increased novelty-induced locomotion and exploration in the OF in a similar but distinct model of selective *Hcrtr1* disruption in DA neurons. These discrepancies may stem from several factors. First, the mouse model used by Xiao *et al* substantially differs from ours in several ways. While DA^VTA^ cells of Xiao *et al*’s mice predominantly express *Hcrtr1*, we showed that >80% of TH^+^ cells in the ventral midbrain of our mice (in a C57BL/6N X C57BL/6J genetic background) express both *Hcrtr1* and *Hcrtr2* genes^22^. Most critically, Xiao *et al* used a different targeted *Dopamine transporter* (*Dat*)-Cre driver allele^60^, which, unlike ours^61^, lacks an IRES, and thus induces a *Dat* gene KO. Xiao *et al*’s DA-*Ox1R*-KO mice are therefore heterozygous for the *Dat* gene, while our DA*^Ox1R-KO^* mice are not. Reduced *Dat* expression is well documented to increase DA levels and induce locomotor hyperactivity^62^ and indeed heterozygous Dat-Cre mice lacking an IRES show locomotor hyperactivity^63^, which could explain some of Xiao *et al*’s observations. In contrast, we use *Dat^+/ires-Cre^* mice reported by Backman *et al* to show unaltered extracellular DA levels^61^, and we moreover reported that *Dat^+/ires-Cre^*mice display normal locomotor activity as *Dat^+/+^* mice^64^. Second, Xiao *et al* used an OF arena measuring 27 × 27 cm, while ours was 45 x 45 cm, thus considerably larger, potentially explaining reduced OF exploration in our case. Strengthening this interpretation, while we found reduced exploration of DA*^Ox1R-KO^* mice in large open fields, we also find that our mice display hyperactivity in more confined arenas.

DA*^Ox1R-KO^* mice also featured potential indicators of impaired risk assessment. These included atypical responding in the EPM trial 2, performed 24 h after trial 1. While mice typically show diminished exploration of open arms in the 2^nd^ EPM trial, as do control mice, DA*^Ox1R-KO^* mice show unabated exploration, suggesting alterations in behavioral adaptation or in acquisition of phobic-like responses^47,48^. Whether apparent lack of behavioral adaptation of DA*^OX1R-KO^* mice in EPM trial 2 reflects altered risk assessment requires further tests. Interpretation of impaired risk-assessment may align with reports by others that OX→NAc^D2^ cell activation is critically involved in innate risk-avoidance in mice^46^, and the discovery that NAc^D2^ cell activity encodes unfavorable outcomes, that drives learned risk-avoidance^65^. DA*^OX1R-KO^* mice nevertheless successfully learnt the active avoidance task, showing as many successful avoidance responses as controls, albeit they exhibited shorter latencies than controls to run to the safe chamber to avoid the shock when exposed to the tone.

Of note, OXs act by postsynaptic but also extrasynaptic action, i.e., on cell somata and dendrites but also at the level of axonal terminals. OX neurons heavily project to the NAc, and OX is known to be able to modulate the release of neurotransmitters, such as glutamate, GABA and DA, from respective axonal terminals^66–69^. Thus, the effects of our DA cell-specific KO mice may be exerted at multiple levels, including by the loss of OXRs on DA terminals, f.i. on accumbal terminals^66,67^. Further studies, notably using local administration of OXR antagonists, are required to distinguish these possibilities.

Altogether, the traits displayed by DA*^Ox1R-KO^* mice, decreased exploration of central zones and the novel object in the OF, hyperactivity in EPM trial 1 or the 3-chamber arena, absence of reduced time in open arms in EPM trial 2, premature and repetitive responding in 3-CSRTT, precipitated avoidance in the active avoidance task, all indicate altered exploration and/or anxiety-like behavior. These changes could be hypothesized to stem from alterations in information processing. Specifically, mesocortical projections (DA^VTA^→PFC) are thought to help calibrate DA levels, so as to optimize signal-to-noise ratios, to enhance discrimination of the most relevant stimuli and facilitate disengagement of motor outputs following decision-making^70–73^. DA*^Ox1R-KO^* mice’ behavioral deficits may represent deficiencies in these pathways, due to inefficient OX_1_R-dependent DA cortical release. Consequent low discrimination of salient stimuli may induce suboptimal goal-oriented and exploratory behavior, as well as activation of inefficient motor responses, as observed in the EPM and 3-CSRTT. Task-locked photometric measures of DA release using DA sensors in multiple DA target fields such as the NAc and mPFC of our mice should shed light on these questions.

### DA*^Ox2R-KO^* mouse behavior: constitutively hyperaroused, strong in reward-driven but compromised in avoidance-driven learning and sociability

Our prior and present data show that dopaminergic *OX_2_R* genetic loss generates a complex phenotype, with a combination of strengths and weaknesses relative to genetic controls. On one hand, mice exhibit a dramatic increase in EEG markers of alertness, enhanced EEG theta-gamma phase-amplitude coupling, markedly improved learning and attentional accuracy in a reward-driven operant learning task^22^. On the other hand, they display poor impulse control^22^ and severely compromised aversion-driven instrumental learning. Hence, DA*^Ox2R-KO^* mice’ electrocortical hyperarousal is linked to advantages in some tasks (attention for positively-valued stimuli) but not others (learning negatively-valued cues), suggesting distinct effector pathways involved in reward *vs* aversive learning.

Higher cognitive performance in reward-driven learning tasks but lower performance in avoidance-driven tasks, or vice versa, is observed in mice. For instance, C57BL/6 mice typically perform well in reward-motivated tasks but show less robust performance in aversive tasks compared to other strains such as DBA/2J, which are known for heightened anxiety-like behavior, fair avoidance learning, but appear less motivated by rewards. These cognitive style differences are sometimes categorized as ‘reward-sensitive’ *vs* ‘punishment-sensitive’ learning phenotypes. Whether a trade-off between reward-sensitivity and avoidance-sensitivity exists is unclear^74,75^.

Enhanced learning and attentional performance in 3-CSRTT could be a response of enhanced DA^VTA^ activity resulting from the constitutive loss of OX_2_R-dependent brake on DA^VTA^ activity, as discussed below. In contrast, failure of DA*^Ox2R-KO^* mice to learn efficiently the active avoidance task may reflect unbalanced DA cell activation, as alteration in DA neuronal activity was documented to disrupt aversive conditioning and trauma-related generalized anxiety-like responses^76^. A specific hypothesis is that impaired active avoidance of DA*^Ox2R-KO^*mice results from altered mesolimbic pathways. D2^NAc^ cells play critical roles in action selection whereby D2^NAc^ excitation governs avoidance of learned risks^65^. Enhanced DA^VTA^ activity in DA*^Ox2R-KO^* mice may lead to increased NAc DA release, and since DA inhibits D2^NAc^ cells, reduced D2^NAc^ cell activity, causing diminished risk-avoidance learning^65^. Blomeley *et al*^46^ moreover showed that a direct OX→D2^NAc^ excitatory circuit is necessary for innate risk-avoidance. Our data suggest that an OX→D2^NAc^ excitatory pathway is potentially also critical for learned risk-avoidance.

Contingency awareness, or learning to link external cues to aversive outcomes, is DA-dependent and a critical factor in determining generalized anxiety-like traits. Ventral striatum voltammetry recordings showed that differences in DA levels are associated with differences in behavioral outcomes in the active avoidance test, whereby DA encodes a safety prediction error signal^77^. In addition to mesolimbic pathways, DA^VTA^→mPFC mesocortical projections also participate in active avoidance learning^78^. Although the identification of the downstream effectors of the effects we observe mandate further work, our data strongly suggest that signaling via OX_2_R, rather than OX_1_R, in DA cells is critical for active avoidance learning.

A major novel salient finding of our study is that dopaminergic OX_2_R loss causes diminished social preference in male mice. OX neurons were previously shown to be involved in social interaction^53,79^. For instance, acute OX cell optogenetic inhibition leads to male mice engaging less in social interactions, in an apparently OX_1_R- and VTA-dependent manner^53^. That study however lacked OX target cell-specificity and thus failed to identify DA cell involvement. Work using whole-body-*Hcrtr1*-KO mice^54^, or systemic OX_1_R antagonism^53^ already implicated OX_1_R in social behavior, but again without a cell type-based mechanistic framework. Our work focusing on OX signaling selectively in DA neurons revealed that loss of OX_2_R, but not OX_1_R, compromises the time male mice engage in social interactions in the 3-chamber social preference task. Future experiments assessing the differential contribution of OX_1_R and OX_2_R signaling in DA neurons in both sexes and additional experimental paradigms are warranted to decode the function of OX signaling in social discrimination.

DA*^Ox2R-KO^* mice overall feature several endophenotypes reminiscent of mouse models of autism spectrum disorder (ASD): enhanced performance in some types of learning but not others, pronounced compulsivity, and compromised social interactions align with behavioral traits seen in ASD mouse models^80^. The use of mice as ASD models, however, remains controversial^81,82^. Nonetheless, since alterations in DA^VTA^→NAc/mPFC pathways are increasingly implicated in ASD-related traits^83,84^, probing these circuits in DA*^Ox2R-KO^*mice for sites of altered DA release may help identify the neurophysiological mechanisms of their behavioral phenotypes and define their relatedness to established ASD mouse models.

Pronounced compulsivity and impulsivity were the only behavioral characteristics common to both DA*^Ox1R-KO^* and DA*^Ox2R-KO^* mice. Therefore, impaired executive control in the form of premature or repetitive responding may represent a point of convergence and major effect of disrupting OX→DA pathways.

### Linking mouse behavioral profiles with *ex vivo* DA^VTA^ cell electrophysiology

The contrasting behavioral phenotypes of DA*^Ox1R-KO^* and DA*^Ox2R-KO^* mice suggest that OX_1_R and OX_2_R receptors contribute differently to DA cell activity and DA synaptic circuits. To assess whether OX_1_R and OX_2_R mediate distinct electrophysiological effects on DA^VTA^ cells, and to isolate cell-autonomous responses independent of network effects implicating non-DA^VTA^ cell types, we first performed patch-clamp recordings of DA cells acutely dissociated from brain VTA slices. In addition, we conducted patch-clamp recordings of DA cells in VTA slices of WT and our mutant lines. To address mesolimbic pathways, we focused of DA neurons of the lateral VTA, which preferentially project to the NAc shell and respond to OXA with increased firing^39^. We aimed at isolating alterations in intrinsic excitability of DA^VTA^ cells in response to OXs. Thus, in VTA slices, we did not assess spontaneous firing but stimulated the cells to fire at regular successive timepoints. Forcing the cells to fire highlights the contribution of altered intrinsic excitability.

In both preparations, we found differential effects of OXA vs OXB, with OXA increasing DA^VTA^ cell firing in WT or controls, but not DA*^Ox1R-KO^* mice. Although OXA is reported to bind equally well OX_1_R and OX_2_R, our data indicate that the DA^VTA^ cell response to OXA is primarily OX_1_R-driven, consistently with prior results showing OX_1_R antagonist SB-33487-mediated blockade of OXA effect on DA^VTA^ cell firing^39^. On the other hand, OXB and the OX_2_R-selective agonist OXB-AL, induced an inhibitory and OX_2_R-dependent response. In DA^VTA^ cells of DA*^Ox2R-KO^* mice, the OXB-mediated firing decrease observed in control mice was prevented, and a tendency towards increased firing was observed, potentially explained by a residual degree of response of OX_1_Rs to OXB. Such interpretation would be consistent with the higher inhibitory effect displayed by OXB-AL, compared to OXB, in dissociated neurons.

These results support the view that OXA-induced OX_1_R signaling stimulates, while OXB-mediated OX_2_R signaling inhibits, DA^VTA^ cell firing. Consequently, deletion of *Hcrtr2* in DA^VTA^ neurons should relieve an inhibitory component of OXR signaling and facilitate OX excitatory effects. This aligns with our observation that DA*^Ox2R-KO^* mice exhibit EEG theta-enriched wakefulness and REMS, and with our hypothesis that theta power increase stems from relief of inhibition of the VTA-septo-hippocampal pathways that generate theta activity^22^. Moreover, our findings that OXA enhances DA^VTA^ neuron firing predominantly via OX_1_R, while OXB decreases firing via OX_2_R suggest that OX_1_R activation promotes reward-related behaviors, while OX_2_R activity may restrain overexcitation to maintain behavioral flexibility.

Although OX_2_R signaling is excitatory in various cell types, such as histaminergic neurons⁷⁸, it is well established that OXR signal transduction mechanisms vary across cell types⁷⁹ and inhibitory effects are not unprecedented. OXR-mediated inhibition has been evidenced in POMC, SCN, and a subset of MCH neurons^85–88^. The intracellular mechanisms by which OX_2_R signaling is intrinsically inhibitory in DA^VTA^ neurons remain to be determined. While OX_1_R has been primarily described to couple to G_αq_ and activate phospholipase C, OX_2_R was shown to couple to both G_αq_ and G_αi/o_-classes of G-proteins^89^. Both pathways can modulate K^+^ currents^90^ and thus potentially cause firing inhibition. G_αq_-dependent Ca^2+^ mobilization from intracellular stores could stimulate Ca^2+^-activated K^+^ channels. Based on evidence in 5HT dorsal raphe nuclei, small conductance Ca^2+^-activated K^+^ (SK/K_Ca_) channels^91^, that are expressed in DA^VTA^ cells^92^, are candidate targets. In this line, recent findings suggest that modulating K_Ca_ currents that inhibit DA^VTA^ firing may regulate autistic-like traits in mouse models^93^. KCNQ (‘M’) K^+^ channels can also be modulated by G_βγ_ subunits^94^. KCNQ are operant in DA^VTA^ neurons and are sensitive targets for behavioral modulation^95^. Finally, although the physiological relevance of G_αi_-mediated pathway downstream of OX_2_R is uncertain⁷⁹, G_iβγ_ subunits are known to activate inwardly rectifying K^+^ channels and this mechanism underlies the inhibitory effect of D_2_ autoreceptors in DA^VTA^ cells^110^.

Our results are in contrast to Korotkova *et al*^32^, reporting increased DA^VTA^ cell firing by both OXA and OXB in VTA slices from 3-4-week-old Wistar rats. These authors, however, recorded spontaneous firing. In such conditions, DA cell firing results from integration of cell-autonomous effects combined with exogenous contributions, such as local GABAergic input as well as terminals from mPFC glutamatergic afferents. Additionally, while our slice recordings targeted putative NAc-projecting neurons, Korotkova *et al*^32^ did not attempt output specificity.

Our findings highlight striking differential effects of the two OX peptides and the two OX receptors on DA^VTA^ cells. Further studies are essential to explore the molecular mechanisms and intracellular targets of these effects, define the differential dynamics and interactions of OX_1_R and OX_2_R signaling, and importantly to assess *in vivo* how OXA and OXB compete to exert their opposing modulation of DA^VTA^ cell activity.

### Limitations

While we use pan-neuronal (*Dat*-driven) mouse models, our electrophysiological analyses focus on DA^VTA^ cells (A10 neurons), as a major and best characterized DA OX target, involved in the behavioral endpoints of interest, and showing major alterations upon loss of OX_1_R or OX_2_R signaling. However, other DA populations may also be involved in our phenotypes, although whether they express *Ox_1_R* and/or *Ox_2_R* is not always known. The responses of nigral DA cells to OX peptides remain controversial^96–98^. Hypothalamic A11-A15 or dorsal raphe nucleus DA cells are implicated in vigilance regulation and behavioral phenotypes^99,100,101^, and could be involved in our mice behavioral profiles. Korchynska *et al*^101^ identified a hypothalamic DA locus implicated in psychostimulant-induced hyperlocomotion in mice. OX_2_R activation in dorsal raphe DA cells may contribute to stress resilience and arousal regulation. A systematic analysis of different DA cell groups to assess their involvement in the behavioral phenotypes we observed is thus warranted.

DA cell heterogeneity in receptor expression, electrophysiological properties and cell connectivity in midbrain ^102^ and other DA nuclei is moreover a confound of all studies using pan-dopaminergic Cre drivers. In the temporal domain, since *OxR* gene inactivation occurs in our mice when *Dat-IRES-Cre* is first expressed, i.e., at around E17^61^, or shortly before birth, developmental compensations may intervene. Of note, OX neurons and innervation essentially mature 3 weeks postnatally in mouse^103^. To ask whether *OxR* gene inactivation in specific DA nuclei of adult mice leads to similar or different phenotypes as those described in the present models, our *OxR^flox/flox^*mice can be injected with *Cre* expressing-vectors in DA nuclei of interest.

Lastly, our behavioral study only followed male mice. In view of the growing evidence of sexual dimorphism in OX- and DA-mediated behaviors^53,104,105^ analysis of female mice is mandated in later studies. Specifically, while both male and female mice exhibit increased locomotion and exploration when DA OXR signaling is disrupted, females may display unique phenotypes, such as increased stereotypic activity^106^. OX activation was reported to be higher in female than male rats and contribute to differences in stress responses and cognitive flexibility, traits that are relevant to neuropsychiatric conditions that disproportionately affect women (Grafe et al., 2017). Inclusion of female mice will enable better understanding of the mechanisms underlying sex-specific vulnerabilities and responses in OX- and DA-dependent disorders.

### Implications for neuropsychiatry

Our data establish a genetically-defined link between monosynaptic OX→DA neurotransmission and disease-relevant socio-emotional behaviors and executive cognition. We evidence an unprecedented dichotomy in dopaminergic OX_1_R *vs* OX_2_R function for DA^VTA^ cell activity and endophenotypes affected in neuropsychiatric disorders including obsessive-compulsive, attention-deficit/hyperactivity, and autism-spectrum disorders. Specifically, our study is the first to report OXB-mediated inhibition of DA^VTA^ cell activity through an OX_2_R-dependent process, as demonstrated by its abrogation in full-body *Hcrtr2-*KO mice (**Fig. 1**), in DA-specific OX_2_R-KO mice (**Fig. 3d-f**), and by OX_2_R-selective antagonism (**Fig. 3g-i**). In parallel, we confirm OX_1_R-mediated enhancement of DA^VTA^ cell excitability by OXA. As the DA system is a primary OX target and is affected in multiple neuropsychiatric disorders and neurological diseases, detailed understanding of OX-mediated neuromodulation of the DA system has great translational relevance.

In the wake of the growing interest in OXR-targeted drug development—whether as single or dual OXR agonists or antagonists for the treatment of a variety of disorders such as substance abuse, eating, obsessive-compulsive, anxiety, pain-, sleep-related disorders and others^37,107–111^ — a thorough understanding of the distinct physiological effects of the two receptors is essential for anticipating potential side effects of these novel therapies.

Dual OXR antagonists, such as Suvorexant, are already used for the treatment of insomnia, and are considered for opiate-addiction^109^. As we show that both DA*^Ox1R-KO^* and DA*^Ox2R-KO^* mice exhibit pronounced compulsivity-impulsivity related endophenotypes, side effects of dual OXR antagonists in this domain should be carefully monitored.

OX_1_R-selective antagonists are gaining interest for use in treatment of substance abuse, binge eating, obsessive, and compulsive disorders. While blocking OX_2_R reduces wakefulness, the expected advantage of selective OX_1_R antagonism may be the ability to achieve clinical efficacy without sleep promotion^112^. As we show that DA OX_1_R ablation induces an anxiety-like effect, OX_1_R antagonists should be evaluated for potential anxiogenic effects.

Our data that OXB and other OX_2_R agonists diminish DA^VTA^ neuronal excitability appeal to a close monitoring of DA-related traits in patients treated with the expected soon to be released OX_2_R-selective agonists for narcolepsy type-1 and obstructive sleep apnea. As D2 agonists that inhibit DA^VTA^ neurons were shown to aggravate cataplexy^113^, aggravation of cataplexy may have been a concern. An 8-week-long phase 2 randomized placebo-controlled trial of the OX_2_R agonist Oveporexton (TAK-861) for narcolepsy type-1 patients however reported positive effects on cataplexy^114,115^, suggesting that the DA system is not a primary target of OX_2_R agonists, or at least not in the contexts so far assessed. If long-term usage of OX_2_R-selective agonists generates anhedonia by suppressed DA^VTA^ cell excitability^116^ remains to be determined. In sum, our findings have extensive implications for the emerging field of OXR neuropharmacology.

## Acknowledgments.

This work was supported by the Swiss National Science Foundation (grant 31003A_182613 to AV), and by the University of Milano-Bicocca (post-doctoral fellowship to LCG, 2021-ATE-0042 and 2024-ATE-0117 to AB). We thank Dr Leonardo Restivo from the NeuroBau Behavioral Facility at the Department of Fundamental Neurosciences of the University of Lausanne for help, notably in the Active Avoidance test, and Anne-Catherine Thomas for animal genotyping.

## Author Contributions

ST conceived, performed and analyzed behavioral experiments with the help of RK and SR for the 3-CSRTT; SA conceived, performed and analyzed patch clamp recordings of DA cells in brain slices; LCG, FG, and AB conceived, performed and analyzed patch clamp recordings of dissociated DA cells; GL participated in study interpretation and schematic design; AV generated the *Hcrtr1* and *Hcrtr2* floxed and whole-body mouse lines; AV, AB, and MT conceived and supervised the project. All authors contributed to manuscript writing and Figure design.

## Competing Interests

Authors declare no competing interests.

## Disclosure

An earlier version of the manuscript was uploaded on BioRxiv. https://www.biorxiv.org/content/10.1101/2025.02.14.638329v1

## SUPPLEMENTAL INFORMATION

## Supplemental Methods and Materials

### Animals

Mice were generated and bred as previously described^22^. Briefly, we engineered the *Hcrtr1* (*Ox1R*) and *Hcrtr2* (*Ox2R*) genes to create Cre-dependent knockout/GFP-reporter floxed alleles: *Hcrtr1^flox^* and *Hcrtr2^flox^* crossed to a Dopamine transporter Cre driver (Dat-IRES-Cre), generating:

*Hcrtr1^flox/flox^;Dat^+/Cre^* (abbreviated: DA*^Ox1R-KO^*) and *Hcrtr1^flox/flox^*;*Dat^+/+^* (DA*^Ox1R-CT^*) controls, and *Hcrtr2^flox/flox^;Dat^+/Cre^* (DA*^Ox2R-KO^*) and *Hcrtr2^flox/flox^* (DA*^Ox2R-CT^*) controls. We therefore generated 4 genotypic groups (2 KO:CT pairs, for *Hcrtr1* and *Hcrtr2*, respectively), and performed all analyses as pair-wise comparisons between KO and CT littermate groups.

Electrophysiological recordings used KO and CT littermate mice that were not subjected to any behavioral test. Behavioral tests used 3- to 6-month-old males that were housed in groups of 2-4 mice/cage and maintained on a 12h:12h light–dark cycle (light-on at 07:00 AM). Mice had *ad libitum* access to food and water, except from cohorts assessed in the 3-Choice Serial Reaction Time Task that were maintained under food restriction, according to the procedure described below. The number of mice in each genotype group is described in detail below in *Behavioral tests*.

### Patch-clamp electrophysiological recording of dissociated DA^VTA^ neurons

Cells were dissociated from 400 μm-thick acute brain slices prepared from P13 to P18 young mice, following standard procedures^117^. Midbrain coronal slices were cut between +3.88 and +2.92 mm from bregma^118^ and left to recover for ≥1 h at 30°C under perfusion with oxygenated solution (95% O_2_, 5% CO_2_), containing (mM) 87 NaCl, 21 NaHCO_3_, 1.25 NaH_2_PO_4_, 7 MgCl_2_, 0.5 CaCl_2_, 2.5 KCl, 25 glucose, 0.4 ascorbic acid, 75 sucrose. The dissociation procedure followed Zhang *et al*^119^, with some modification. Slices were incubated in a medium (DM/KYN) containing (mM): 134 Na-isethionate, 23 glucose, 2 KCl, 4 MgCl_2_, 1 CaCl_2_, 1 kynurenic acid, 15 HEPES, and supplemented with 0.75 mg/ml protease type XIV (Sigma P-5147). Enzymatic treatment was carried out at 37°C, for a time proportional to mice age (from 19 min for P13, to 24 min for P18). After rinsing with DM/KYN, the VTA region was cut out with a polished needle under a stereomicroscope and laid on a Petri dish pretreated with concanavalin A (50 μg/ml). After delicate mechanical dissociation with a fire-polished micro-Pasteur pipette, cells were left at 30 °C in a humidified chamber, for at least 30 min. DA cells were identified by their slow spontaneous discharge frequency (2-3 Hz on average^32,120^ and the rapid blockade of cell firing by 100 μM DA, which activates inward rectifying K^+^ channels by engaging D_2_ autoreceptors^121^. Applying DA before or after testing the OX response did not alter cellular response, thus ruling out potential cross talk between D_2_Rs and OXRs. Patch-clamp recordings were carried out in cell-attached configuration at room temperature using an Axopatch 200B amplifier (Molecular Devices). Signals were low-pass filtered at 2 kHz and digitized at 10 kHz, with pClamp9/Digidata 1322A (Molecular Devices). During the experiments, neurons were perfused with an extracellular solution containing (mM): 145 NaCl, 1 MgCl_2_, 3.5 KCl, 2 CaCl2, 5 glucose, 10 HEPES (pH 7.25). The pipette contained (mM): 140 K-gluconate, 10 NaCl, 1 MgCl_2_, 2 MgATP, 0.3 NaGTP, 0.1 BAPTA, 10 HEPES (pH 7.3). Drugs were applied with an RSC-160 Rapid Solution Changer (BioLogic Science Instruments). OX peptides were applied for 2 min, after at least 3 min of baseline (Control) recording. Recovery was generally followed for approximately 10 min (Fig. 1 reports the firing frequency for the first 2 min after peptide washout).

### Patch-clamp electrophysiological recordings of DA neurons in VTA brain slices

Mice (10- to 13-week-old) were anaesthetized with isoflurane and decapitated. The brain was quickly removed and coronal slices (200 µm-thick) containing the midbrain were prepared using a vibrating tissue slicer (Campden Instruments, Loughborough, UK) in oxygenated (95% O_2_ / 5% CO_2_) ice-cold modified artificial cerebrospinal fluid (aCSF) containing (in mM): 105 sucrose, 65 NaCl, 25 NaHCO_3_, 2.5 KCl, 1.25 NaH_2_PO_4_, 7 MgCl_2_, 0.5 CaCl_2_, 25 glucose, 1.7 L(+)-ascorbic acid. Slices recovered for 1 h at 35°C in standard aCSF containing (in mM): 130 NaCl, 25 NaHCO_3_, 2.5 KCl, 1.25 NaH_2_PO_4_, 1.2 MgCl_2_, 2 CaCl_2_, 18 glucose, 1.7 L(+)-ascorbic acid, and complemented with 2 sodium pyruvate and 3 myo-inositol. In the recording chamber, slices were superfused with oxygenated standard aCSF at nearly physiological temperature (30-32°C). In all patch-clamp recordings of DA cells from VTA slices, we targeted lateral VTA cells, reported to project to the lateral NAc-shell, based on their location and presence of an I_h_ current^39^. Neurons were patched in the whole-cell configuration with borosilicate glass pipettes (2-4 MΩ) filled with (in mM): 130 K-gluconate, 10 KCl, 10 HEPES, 10 Na-phosphocreatine, 0.2 EGTA, 4 Mg-ATP, 0.2 Na-GTP, (290-300 mOsm, pH 7.2-7.3). Recorded neurons were considered dopaminergic if they exhibited an inward current/sag in response to hyperpolarizing voltage/current steps, as well as a >3 mV hyperpolarization upon perfusion of the GABA_B_R agonist baclofen (1 µM) at the end of the recording^122^. To assess basal intrinsic excitability, cell firing was first elicited by providing 1-s long incremental somatic current injections while the membrane potential was held at -60 mV. To study the effect of OXA and OXB on intrinsic excitability, we used a protocol previously described^39^. Briefly, neuronal firing was elicited by a depolarizing current injection of 500 ms duration and of constant amplitude. Following a baseline period of 10 min, 100 nM OXA or OXB was bath-applied for 5 min. Throughout these recordings, cells were held at -60 mV by direct current injection, in order to minimize the contribution of synaptic depolarization/hyperpolarization and isolate the effect of orexin peptides on intrinsic excitability. Baimel *et al*^39^ demonstrated that the effect of OXA is independent of synaptic transmission. We also verified that the effect of OXB on DA^VTA^ cells persists in the presence of synaptic blockers (**Fig. 7h-i**) (100 µM picrotoxin, 10 µM DNQX and 50 µM D,L-APV, to antagonize GABA_A_Rs, AMPARs and NMDARs, respectively).

Recordings in the current-clamp configuration were conducted with bridge balance. Membrane voltage values were not corrected for liquid junction potential. Data were acquired through a Digidata1550A digitizer. Signals were amplified through a Multiclamp700B amplifier (Molecular Devices, Sunnyvale, USA), sampled at 20 kHz and filtered at 10 kHz using Clampex10 (Molecular Devices). Clampfit10 (Molecular Devices) was used for data analysis.

### Behavioral tests

#### Behavioral pipeline

Mice (11 DA*^Ox1R-KO^* and 11 DA*^Ox1R-CT^*; 12 DA*^Ox2R-KO^* and 19 DA*^Ox2R-CT^*) performed sequentially the open field test, the elevated plus maze test, the 3-chamber sociability test and the active avoidance test, with ≥24 h between each test. Active avoidance was assessed last as it entails exposure to electric shocks, hence is the most stressful. A separate cohort of 7 DA*^Ox1R-CT^* and 8 DA*^Ox1R-CT^* performed the 3-choice serial reaction time task (3-CSRTT). Rationale for animal exclusion from analysis is given in detail for each individual behavioral test below.

#### Open field (OF) and reactivity to novelty (NO) tests

To assess locomotion, exploration, and anxiety-like behavior, an open field test adapted from previous studies^91,92,123^ was performed, using a Plexiglas (Length x Width x Height) 45 x 45 x 39.5 cm arena equipped with a ceiling-mounted camera. The test consisted of 2 parts: (1) the mouse is allowed to freely explore the open field for 10 min; (2) as the mouse stays in the open field arena, a small object is then softly brought from the side and placed in the arena center, and the mouse allowed to freely explore the arena and the object for a further 5 min, during which reactivity to the novel object is assessed. The arena was divided in center and wall zones, while for the Novel Object Reactivity test, the center zone was further divided to contain a smaller zone around the object (15 x 15 cm). The behavior was monitored using the video camera and analyzed with a computerized tracking system (Ethovision XT 15, Noldus IT). Entries and time spent in the exploration area (center) and the periphery (wall), as well as the total distance travelled in the arena, were recorded automatically. The light intensity in the arena was adjusted to 12-15 lux. The apparatus was cleaned with 5% Ethanol solution and dried thoroughly after each animal. One mouse out of 11 from the DA*^Ox1R-KO^* group was excluded from this test as it climbed on the object for a prolonged time.

#### Elevated plus maze

To assess anxiety-like behavior, mice were subjected to the elevated plus maze (EPM) test according to previously published protocols^123,124,93,94^. The elevated plus maze consists of a cross-shaped arena with two open arms and two closed arms with walls made of Plexiglas and an open center (80 x 78 x 15 cm), which is elevated 47 cm above ground. In each trial, the mouse is placed in the center facing a closed arm and allowed to freely explore the maze for 5 min. Behavior is monitored using a video camera and analyzed using a computerized tracking system (Ethovision XT 15, Noldus IT). Time spent in the center, open and closed arms, and number of visits (entries) into different arms were recorded automatically. The light intensity was 15-16 lux in the open arms, 5-6 lux in the closed arms and 10-12 lux in the center. The apparatus was cleaned with 5% ethanol solution and dried thoroughly after each animal. One mouse out of 11 from the DA*^Ox1R-CT^*group was identified as an outlier and excluded from this test.

#### Three-chamber sociability test

We used a social interaction assay comprising a rectangular Plexiglas arena (60 × 41 × 22 cm; Ugo Basile, Varese, Italy) divided into three chambers (each 20 × 41 × 22 (height) cm). The center chamber has doors that can be lifted so the mouse has free access to all chambers. The social preference test was performed as previously^55,125,126,73^. Briefly, each mouse is placed in the arena for a 10 min habituation period, when it is allowed to freely explore the empty arena. After the habituation, 2 fenced enclosures with metal vertical bars (16 cm × 9 cm) are placed in the center of the 2 outer chambers. One enclosure is empty (serving as an inanimate object) whereas the other contains a social stimulus (an unfamiliar juvenile mouse 25 ± 1 day old). The enclosure allows visual, auditory, olfactory, and tactile contact between the experimental mouse and the juvenile. Juvenile mice had been habituated daily to the apparatus and the enclosures for a brief period of time in the 3 days preceding the experiment. During the test, the experimental mouse was allowed to freely explore apparatus and enclosures for 10 min. The position of the empty or juvenile-containing enclosures alternated and was counterbalanced for each trial to avoid bias effects. Sessions were video-tracked using Ethovision XT (Noldus, Wageningen, Netherlands), providing automated recording of time spent around either enclosure using virtual zones designed around them, total distance travelled, and mouse velocity. The social preference score is defined as the time ratio: social/(social+empty). The arena was cleaned with 5% ethanol and dried between trials. In the DA*^Ox1R-CT^* group, 1 of 11 mice was identified as an outlier (ROUT method) and excluded from the social preference analysis. In the DA*^Ox2R-KO^* group, all 12 mice were included, but 1 of 12 mice did not socially interact (social bout number was 0), thus we were unable to calculate the social interaction bout duration for this mouse.

#### 3-Choice serial reaction time task (3-CSRTT)

The 3-CSRTT test assesses continuous attention performance in rodents. We used an operant chamber (14.5 x 22 x 18 cm) made of two opposing smooth dark-grey aluminum walls and two opposing transparent Plexiglas walls equipped with iSpy camera and POLY software (Imetronic, Pessac, France). We used a protocol, including shaping, training, baseline and impulsivity assessment, adapted from prior studies^49,127–130^. The mouse has to learn to nosepoke in one of three LED-lit apertures in the time during which the light is on (correct response). Each aperture’s light is lit in a pseudo-random manner (the sequence of nosepoke activation appears to be statistically random but relying on a deterministic algorithm; a truly random process could result, for example, in one aperture being active 2,3 or more times in a row, thus potentially creating response biases). Each correct response is followed by delivery of a sucrose pellet (Dustless Precision Pellets® 20 mg, Rodent Purified Diet Chocolate Flavor, Bio-Serv) in the food magazine located on the opposite wall of the chamber. If the mouse does not respond within the timeframe when the light is lit in one aperture, a ‘limited hold’ (LHD) time period follows, during which the LED light is turned off, but the mouse can still perform a correct or incorrect response. An incorrect response, or an omission to respond when the light is on or during the LHD period, leads to a 5-s time-out (TO) period, when all lights are turned off and no action can be taken. Mice were 14 weeks old at the start of the experiment. During 2 weeks, they were habituated to the experimenters with handling and body weight assessment. Mice were then food-restricted to gradually reach 85% of their initial weight, while respecting the expected weight gain for their age. Specifically, we established a mean weekly weight gain of 0.74 g. Weight was monitored every day or every other day. Once the experiment started and mice started to learn to obtain food pellets, we adjusted their regular chow by subtracting the weight of the pellets consumed during the training sessions.

##### Shaping stages

In parallel to food restriction, a ‘Shaping 1’ phase of chamber habituation started when the mice became familiar with the chamber and task contingencies. During this phase, mice received a food pellet reward when they nosepoked in any of the 3 simultaneously lit apertures. To encourage nosepoking, the stimulus lasted 60 s and there were no incorrect responses or TO periods. To go to the 2^nd^ shaping phase (Shaping 2), the mice had to receive ≥20 pellets. During Shaping 2, the mice had to visit the food magazine to initiate a new trial. The stimulus duration was now 30 s and there were still no TO periods after incorrect responses. Unlike in Shaping 1, in Shaping 2, a correct response led to a 2-s ITI before one of the holes was lit again. In case of omissions, the mouse had to start the trial by visiting the food magazine. The mouse had to receive ≥20 pellets to go to the first training stage (i.e. Stage 1).

##### Training stages

Each training session lasted 30 min or 100 trials, whichever condition was met first. The procedure followed the steps and sequence described previously^49^. To start a trial, the mouse has to visit the food magazine and then, an ITI of 2 or 5 s (depending on the stage) is presented before one of the apertures is lit. Premature responses (nosepokes during the ITI), incorrect responses, or omissions led to a 5-s TO and has to be followed by a poke in the food magazine to start the next ITI and trial. Correct responses led to the distribution of a pellet, which led directly to the ITI. Training followed 10 different training stages of gradually increasing difficulties according to **Table S1**. Once performance was as stable as possible in the last training stage, a so-called ‘baseline’ performance for each mouse was determined. One animal from the DA*^Ox1R-KO^* group could not be included in 3-CSRTT analyses as it performed 0 correct responses and showed no learning after 30 days of training.

#### Active avoidance test

To evaluate associative aversion-based learning, we performed an active avoidance paradigm in a 2-way shuttle box (Ugo Basile, Italy). Mice learn that a tone (conditioned stimulus; CS, or ‘warning signal’) precedes a mild electric shock that they can stop by escaping to the adjacent ‘safe’ chamber^131–133^. Mice were run through 50 trials per day for 2 days. Each trial started with a variable interval (15-45 s) after which a 10-s tone preceded a 0.3 mA electric shock delivered through the floor. The shock stopped immediately when the mouse transitioned to the other chamber or after 5 s. The following readouts were quantified: number of premature responses, defined as transitions to the adjacent chamber during the inter-trial interval (ITI) before the tone had started, number of ‘avoidances’, defined as transitions to the safe chamber during the tone (i.e., before the shock), and the number of ‘escapes’, defined as transitions to the safe chamber after the shock had started. Avoidance responses are used as index of learning as they require the mouse to make an association between the CS (tone) and the electric shock to avoid punishment. An automatic system registered avoidances, escapes and premature responses. Latencies were measured automatically by the system as the reaction times to avoidance, premature and escape responses. Non-learners were defined as animals performing fewer than 10 avoidance responses in total across the 2 training days. Between each trial, the apparatus was cleaned with 5% ethanol and dried.

### Statistics

The data were analysed by independent samples *t*-tests, paired samples *t*-tests and repeated-measures ANOVA, as appropriate, using the statistical packages SPSS (Chicago, IL, USA), GraphPad Prism (San Diego, CA, USA), or OriginPro 2023 (OriginLab Corporation). Shapiro-Wilk analysis was used to assess the normality of sample distribution. When normality was violated, non-parametric tests (Wilcoxon or Mann-Whitney, as appropriate) were applied as stated in figure legends. For repeated-measures two-way ANOVA, *P* values of the main effects and interaction are reported in figure legends for each experiment. All bars and error bars represent mean ± S.E.M. Statistical significance was set at *P*<0.05. *P* values were encoded in Figures as \**P*<0.05, \*\**P*< 0.01, \*\*\**P*<0.001. *P*-value was considered trending toward significance when 0.05⩽*P*⩽0.1.

## Illustration software

Figures 4, 5a, 6a, 7a, 8a were created in part with the help of BioRender.com. All Figures were prepared using Affinity Designer 2.

## Data availability

The datasets acquired for this study are available from the corresponding author upon request.

## Code availability

Custom-written scripts for data analysis used in this study are available from the corresponding author upon request

## Supplemental Table

**Table S1.** Criteria defining the test contingencies of the 10 training stages of gradually increasing difficulties used in the 3-choice serial reaction time task test (3-CSRTT) described in Fig. 6. ITI, inter-trial-interval, LHD, limited-hold period.

| Training stage | Stimulus duration (s) | ITI (s) | LHD (s) | Criterion to move to next stage |
| --- | --- | --- | --- | --- |
| 1 | 30 | 2 | 30 | ≥ 30 Correct trials |
| 2 | 15 | 2 | 15 | ≥ 30 Correct trials |
| 3 | 10 | 5 | 10 | ≥ 50 Correct trials |
| 4 | 7 | 5 | 7 | ≥ 50 Correct trials<br>> 70% Accuracy |
| 5 | 5 | 5 | 5 | ≥ 50 Correct trials<br>> 70% Accuracy<br>< 20% Omissions |
| 6 | 4 | 5 | 4 | ≥ 50 Correct trials<br>> 70% Accuracy<br>< 20% Omissions |
| 7 | 3 | 5 | 3 | ≥ 50 Correct trials<br>> 70% Accuracy<br>< 20% Omissions |
| 8 | 2 | 5 | 2 | ≥ 50 Correct trials<br>> 70% Accuracy<br>< 20% Omissions |
| 9 | 1.5 | 5 | 1.5 | ≥ 50 Correct trials<br>> 70% Accuracy<br>< 20% Omissions |
| 10 | 1.2 | 5 | 1.2 | ≥ 50 Correct trials<br>> 70% Accuracy<br>< 20% Omissions |

## Supplemental Figures S1-S4

**Figure S1.**
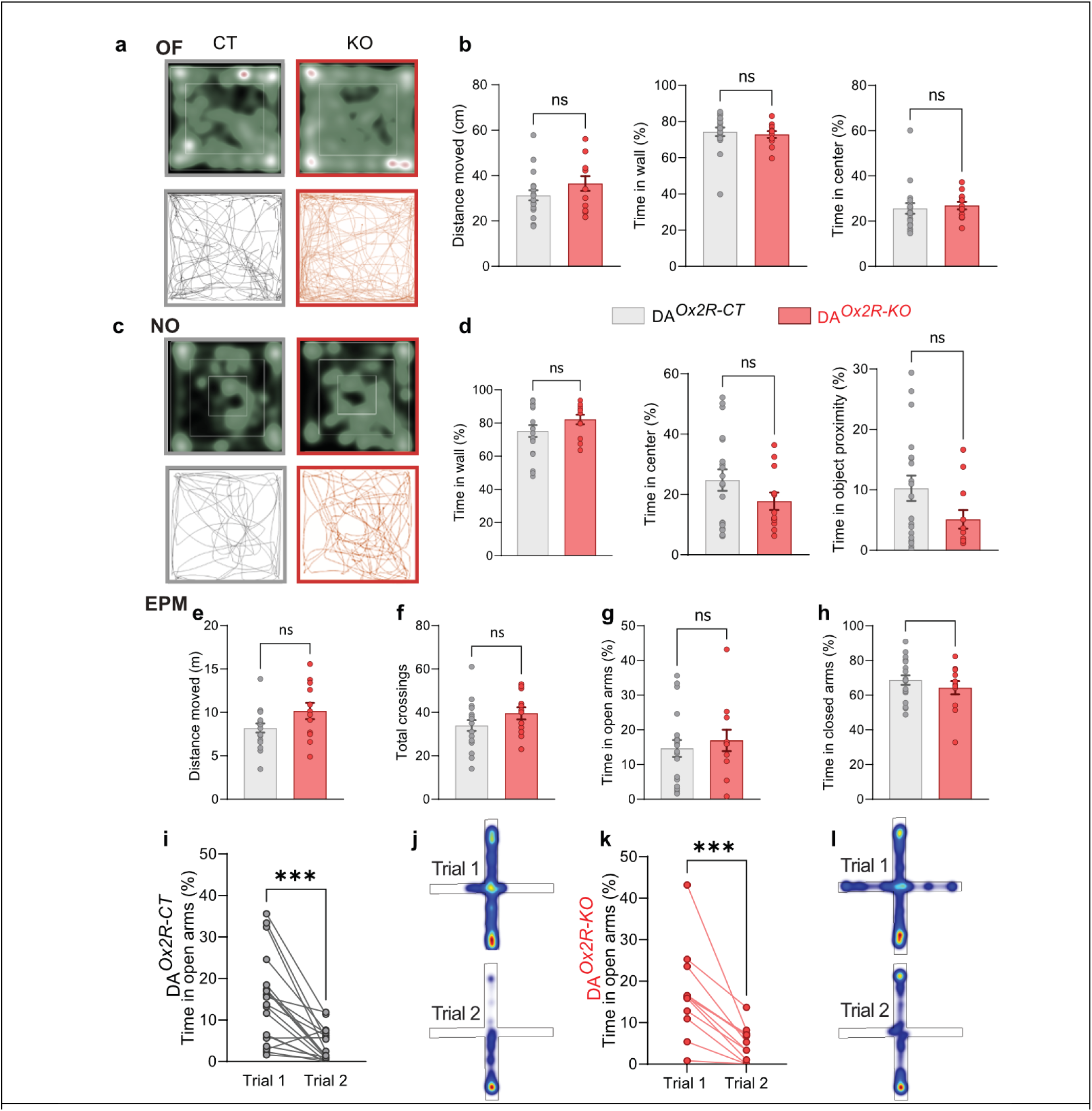
(related to Figure 5). Dopaminergic *Hcrtr2*-ablated mice show normal explorative behavior, reactivity to novelty, and responses in the elevated plus maze. Behavioral profiling of DA*^Ox2R-KO^* and DA*^Ox2R-CT^* mice in the open field (OF), reactivity to novel object (NO) and elevated plus maze (EPM) tests. (**a**) Example heat maps (top) and trail maps (bottom) from DA*^Ox2R-CT^* (left) and DA*^Ox2R-KO^* (right) in the open field arena during 10-min free exploration. (**b**) (Left) distance moved comparison between DA*^Ox2R-CT^* and DA*^Ox2R-KO^* in the open field for 10 min, *t* (29) =1.366, *P =* 0.1883, t-test, middle: comparison between DA*^Ox2R-CT^* and DA*^Ox2R-KO^* for the time in the wall zone for 10 min, *U* = 85, *P =* 0.252, Mann-Whitney test, right: time in the center zone for 10 min in DA*^Ox2R-CT^* and DA*^Ox2R-KO^* (*U* = 86, *P =* 0.269, Mann-Whitney test). (**c**) Example heat maps (Top) and trail maps (Bottom) from DA*^Ox2R-CT^* (Left) and DA*^Ox2R-KO^* (Right) in the arena for 5 min after placing a novel object. (**d**) (Left) comparison between DA*^Ox2R-CT^* and DA*^Ox2R-KO^* for the time in the wall zone during the reactivity to the novel object phase, *t* (29) = 1.383, *P =* 0.177, *t*-test, middle: comparison between DA*^Ox2R-CT^* and DA*^Ox2R-KO^* for the time in the center zone for 5 min during the reactivity to the novel object phase, *t* (29) = 1.389, *P =* 0.175, *t*-test, right: comparison between DA*^Ox2R-CT^* and DA*^Ox2R-KO^* for the time in the center zone around the novel object, *U* = 77, *P =* 0.138, Mann-Whitney test. (**e**) Total distance traveled by DA*^Ox2R-CT^* and DA*^Ox2R-KO^* mice in the EPM, *t* (29) = 1.980, *P =* 0.057, t-test. (**f**) Number of total inter-zone crossings in DA*^Ox2R-CT^* and DA*^Ox2R-KO^* mice, *t* (29) = 1.476, *P =* 0.151, t-test. (**g**) Time in the open arms of the EPM by DA*^Ox2R-CT^* and DA*^Ox2R-KO^* mice (*t* (29) = 0.5586, *P =* 0.562, t-test. (**h**) Time in the closed arms of the EPM by DA*^Ox2R-CT^* and DA*^Ox2R-KO^*, *t* (29) = 0.975, *P =* 0.337, t-test. (**i**) Comparison of the time in open arms between trial 1 and trial 2 for DA*^Ox2R-CT^*, and example heatmaps for trial 1 and trial 2, *W* = -176, *P <* 0.001, Wilcoxon test. (**k**) Comparison of the time in open arms between trial 1 and trial 2 for DA*^Ox2R-KO^*, and example heatmaps for trial 1 and trial 2, *t* (12) = 5.054, *P <* 0.001, paired *t*-test. Bar graphs show individual values and mean ± SEM. *n* = 19 DA*^Ox2R-CT^*, 12 DA*^Ox2R-KO^*.

**Figure S2.**
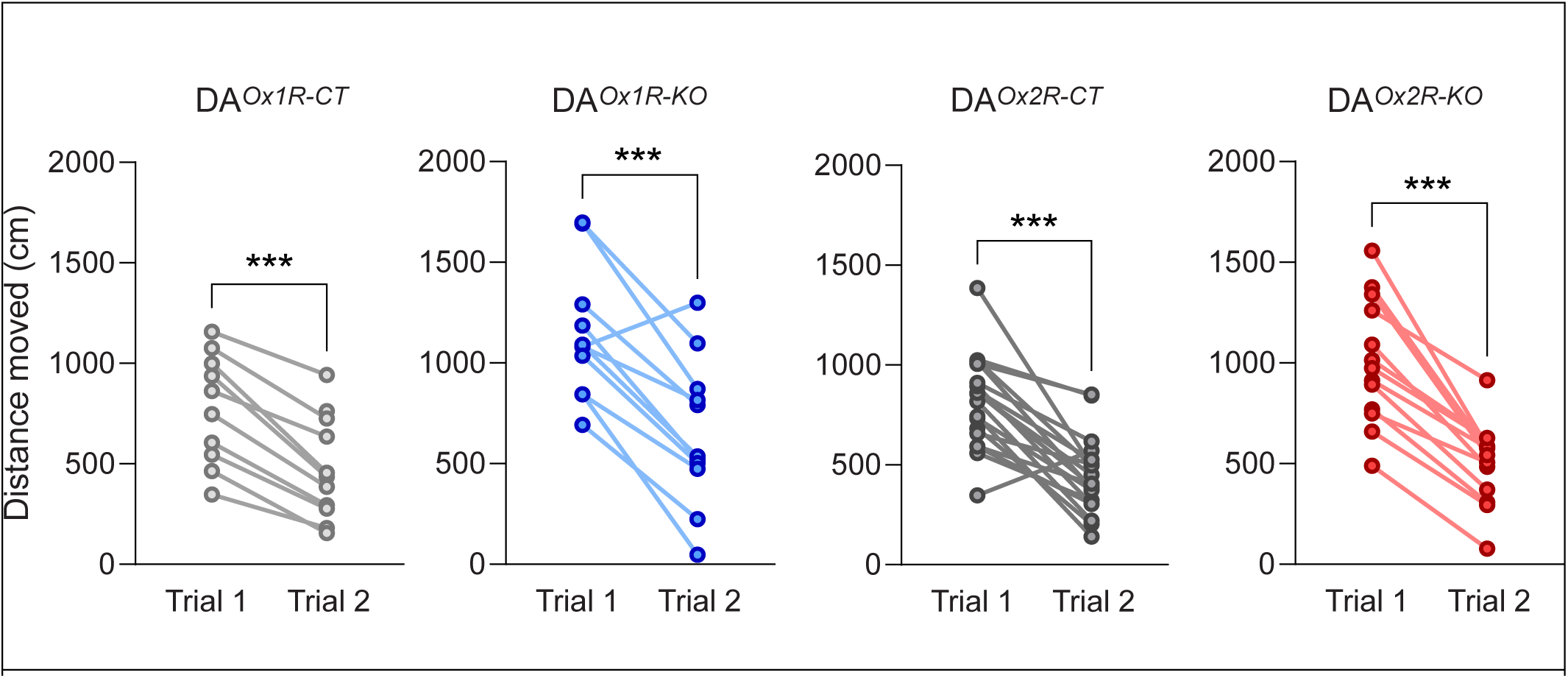
**(related to Figure 5).** Reduced total distance traveled in EPM Trial 2 *vs* EPM Trial 1 in all four genotype groups. Upon re-exposure to a 2^nd^ EPM trial 24 h after the 1^st^ EPM trial, mice of all 4 genotype groups (DA*^Ox1R-CT^*, DA*^Ox1R-KO^*, DA*^Ox2R-CT^*, and DA*^Ox2R-KO^*) exhibit reduced total distance travelled. Difference in distance travelled by the mice of each genotype between the 1^st^ and 2^nd^ trials (paired sample t-test; DA*^Ox1R-CT^*: t (9) = 8.518, *P* <0.001; DA*^Ox1R-KO^*: t(10) = 5.628, *P* <0.001; DA*^Ox2R-CT^*: t (18) = 6.694,*P* <0.001; DA*^Ox2R-KO^*: t(12) = 8.241, *P* <0.001).

**Figure S3.**
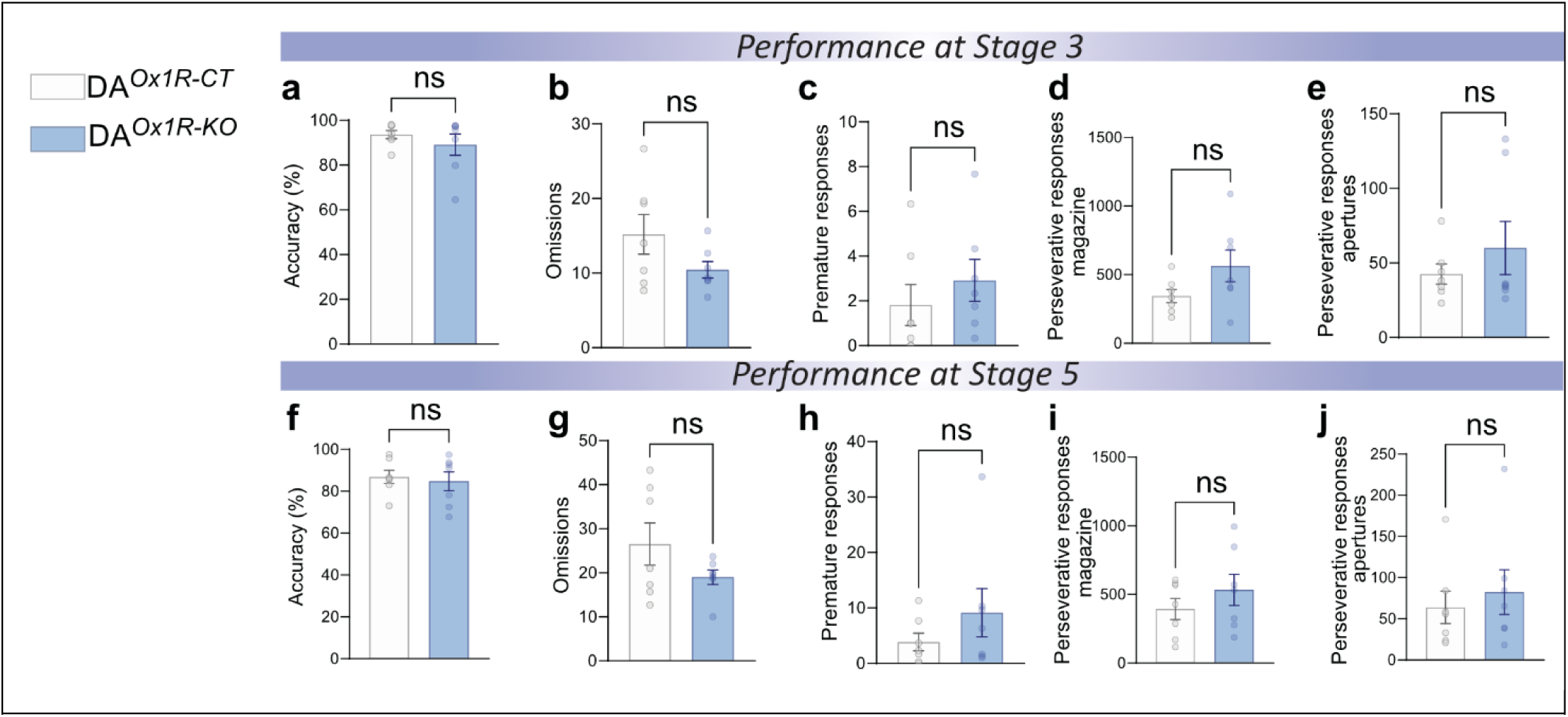
**(related to Figure 6).** Dopaminergic *Hcrtr1*-ablated mice perform similarly as controls in reward-driven operant learning (3-CSRTT) under Stages 3 and 5 training contingencies. (a) Accuracy score at stage 3 in DA*^Ox1R-CT^* and DA*^Ox1R-KO^* mice (*U* = 19, *P =* 0.535, Mann-Whitney test). (b) Omission rate at stage 3 in DA*^Ox1R-CT^* and DA*^Ox1R-KO^* mice (*t* (12) = 1.659, *P =* 0.123, *t*-test). (c) Premature responses at stage 3 in DA*^Ox1R-CT^* and DA*^Ox1R-KO^* mice f(*U* = 14.50, *P =* 0.223, Mann-Whitney test) (d) Rate of perseverative responding in the food magazine at stage 3 in DA*^Ox1R-CT^* and DA*^Ox1R-KO^* mice (*t* (12) = 1.756, *P =* 0.105, *t*-test). (e) Rate of perseverative responding in nosepoking apertures at stage 3 in DA*^Ox1R-CT^* and DA*^Ox1R-KO^* mice (*U* = 23.50, *P =* 0.929, Mann-Whitney test). (f) Accuracy scores at stage 5 of DA*^Ox1R-CT^* and DA*^Ox1R-KO^* mice (*t (12)* = 0.383, *P =* 0.709, *t*-test). (g) Omission responding rate at stage 5 inDA*^Ox1R-CT^* and DA*^Ox1R-KO^* mice (*t* (12) = 1.483, *P =* 0.164, *t*-test. (h) Rate of premature responding at stage 5 in DA*^Ox1R-CT^* and DA*^Ox1R-KO^* mice (*U* = 18.50, *P =* 0.476, Mann-Whitney test). (i) Rate of perseverative responding in the food magazine at stage 5 in DA*^Ox1R-CT^* and DA*^Ox1R-KO^* mice (*t* (12) = 1.015, *P =* 0.330, *t*-test). (j) Rate of perseverative responding in nosepoking apertures at stage 5 in DA*^Ox1R-CT^* and DA*^Ox1R-KO^* mice (*U* = 19.50, *P =* 0.558, Mann-Whitney test). Bar graphs show individual values as mean ± SEM. *n* = 7 DA*^Ox1R-CT^*, 7 DA*^Ox1R-KO^*.

**Figure S4.**
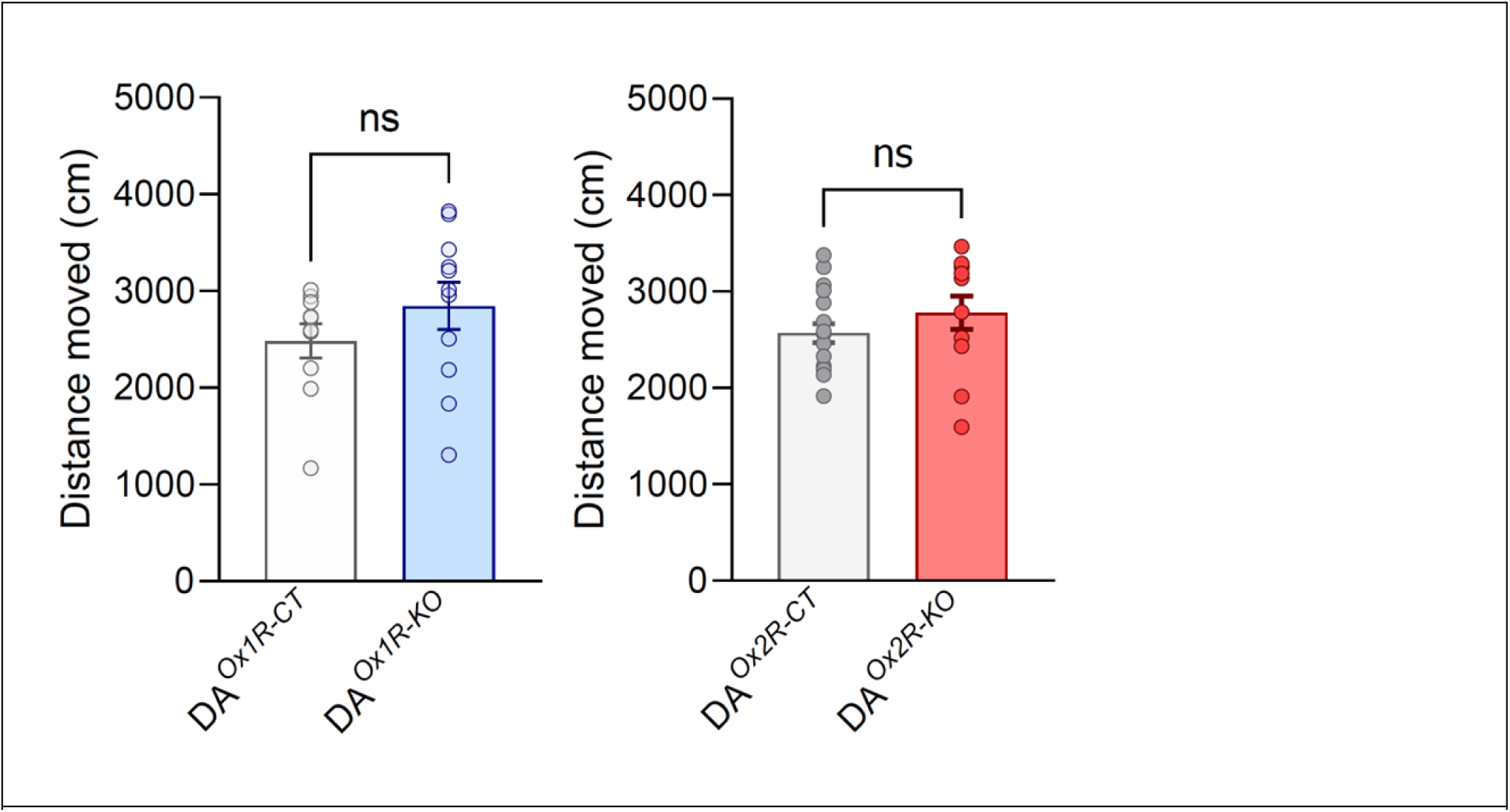
**(related to Figure 8).** DA*^Ox1R-KO^* and DA*^Ox2R-KO^* mice travel across a similar total distance during the social preference phase of the three-chamber test relative to their respective controls (t-test; DA*^Ox1R-CT^* vs DA*^Ox1R-KO^*: t (19) = 1.186, *P* = 0.250; DA*^Ox2R-CT^* vs DA*^Ox2R-KO^* : t (29) = 1.149, *P* = 0.260).

## Notes

### Competing Interest Statement

The authors have declared no competing interest.

### Summary of Updates

Reorganization of the narrative, deeper referencing of the relevant literature; addition of 2 main Figures, including a Summary/Perspective Graphical Abstract Figure. Addition of Author Galina Limorenko for her contribution to conceptual interpretation and Graphical Abstract.

## REFERENCES

1. Mahler SV, Moorman DE, Smith RJ, James MH, Aston-Jones G. Motivational activation: a unifying hypothesis of orexin/hypocretin function. Nat Neurosci 2014; 17(10): 1298–1303.

2. Fronczek R, Lammers GJ, Balesar R, Unmehopa UA, Swaab DF. The number of hypothalamic hypocretin (orexin) neurons is not affected in Prader-Willi syndrome. Journal of Clinical Endocrinology & Metabolism 2005; 90(9): 5466–5470.

3. De La Herran-Arita AK, Zomosa-Signoret VC, Millan-Aldaco DA, Palomero-Rivero M, Guerra-Crespo M, Drucker-Colin R, Vidaltamayo R. Aspects of the narcolepsy-cataplexy syndrome in O/E3-null mutant mice. Neuroscience 2011; 183: 134–143.

4. de Lecea L, Kilduff TS, Peyron C, Gao X, Foye PE, Danielson PE, Fukuhara C, Battenberg EL, Gautvik VT, Bartlett FS, 2nd, Frankel WN, van den Pol AN, Bloom FE, Gautvik KM, Sutcliffe JG. The hypocretins: hypothalamus-specific peptides with neuroexcitatory activity. Proc Natl Acad Sci U S A 1998; 95(1): 322–327.

5. Sakurai T, Amemiya A, Ishii M, Matsuzaki I, Chemelli RM, Tanaka H, Williams SC, Richarson JA, Kozlowski GP, Wilson S, Arch JR, Buckingham RE, Haynes AC, Carr SA, Annan RS, McNulty DE, Liu WS, Terrett JA, Elshourbagy NA, Bergsma DJ, Yanagisawa M. Orexins and orexin receptors: a family of hypothalamic neuropeptides and G protein-coupled receptors that regulate feeding behavior. Cell 1998; 92(5): 1 page following 696.

6. Johnson PL, Truitt W, Fitz SD, Minick PE, Dietrich A, Sanghani S, Traskman-Bendz L, Goddard AW, Brundin L, Shekhar A. A key role for orexin in panic anxiety. Nature Medicine 2010; 16(1): 111–U149.

7. Peleg-Raibstein D, Burdakov D. Do orexin/hypocretin neurons signal stress or reward? Peptides 2021; 145: 170629.

8. Peyron C, Tighe DK, van den Pol AN, de Lecea L, Heller HC, Sutcliffe JG, Kilduff TS. Neurons containing hypocretin (orexin) project to multiple neuronal systems. J Neurosci 1998; 18(23): 9996–10015.

9. Li SB, Damonte VM, Chen C, Wang GX, Kebschull JM, Yamaguchi H, Bian WJ, Purmann C, Pattni R, Urban AE, Mourrain P, Kauer JA, Scherrer G, de Lecea L. Hyperexcitable arousal circuits drive sleep instability during aging. Science 2022; 375(6583): eabh3021.

10. Lee MG, Hassani OK, Jones BE. Discharge of identified orexin/hypocretin neurons across the sleep-waking cycle. J Neurosci 2005; 25(28): 6716–6720.

11. Mochizuki T, Crocker A, McCormack S, Yanagisawa M, Sakurai T, Scammell TE. Behavioral state instability in orexin knock-out mice. J Neurosci 2004; 24(28): 6291–6300.

12. Adamantidis AR, Zhang F, Aravanis AM, Deisseroth K, de Lecea L. Neural substrates of awakening probed with optogenetic control of hypocretin neurons. Nature 2007; 450(7168): 420–424.

13. Broglia G, Bouvier PHP, Tafti M, Bandarabadi M. Orexin mediates neuromodulation during sleep. Journal of Sleep Research 2022; 31.

14. Vassalli A, Franken P. Hypocretin (orexin) is critical in sustaining theta/gamma-rich waking behaviors that drive sleep need. Proc Natl Acad Sci U S A 2017; 114(27): E5464–E5473.

15. Liblau RS, Vassalli A, Seifinejad A, Tafti M. Hypocretin (orexin) biology and the pathophysiology of narcolepsy with cataplexy. Lancet Neurol 2015; 14(3): 318–328.

16. Scammell TE, Winrow CJ. Orexin receptors: pharmacology and therapeutic opportunities. Annu Rev Pharmacol Toxicol 2011; 51: 243–266.

17. Seifinejad A, Vassalli A, Tafti M. Neurobiology of cataplexy. Sleep Med Rev 2021; 60: 101546.

18. Eban-Rothschild A, Appelbaum L, de Lecea L. Neuronal Mechanisms for Sleep/Wake Regulation and Modulatory Drive. Neuropsychopharmacology 2018; 43(5): 937–952.

19. Villano I, Messina A, Valenzano A, Moscatelli F, Esposito T, Monda V, Esposito M, Precenzano F, Carotenuto M, Viggiano A, Chieffi S, Cibelli G, Monda M, Messina G. Basal Forebrain Cholinergic System and Orexin Neurons: Effects on Attention. Front Behav Neurosci 2017; 11: 10.

20. Burgess CR, Scammell TE. Narcolepsy: neural mechanisms of sleepiness and cataplexy. J Neurosci 2012; 32(36): 12305–12311.

21. Okura M, Fujiki N, Kita I, Honda K, Yoshida Y, Mignot E, Nishino S. The roles of midbrain and diencephalic dopamine cell groups in the regulation of cataplexy in narcoleptic Dobermans. Neurobiol Dis 2004; 16(1): 274–282.

22. Bandarabadi M, Li S, Aeschlimann L, Colombo G, Tzanoulinou S, Tafti M, Becchetti A, Boutrel B, Vassalli A. Inactivation of hypocretin receptor-2 signaling in dopaminergic neurons induces hyperarousal and enhanced cognition but impaired inhibitory control. Mol Psychiatry 2023.

23. Mohammadkhani A, Mitchell C, James MH, Borgland SL, Dayas CV. Contribution of hypothalamic orexin (hypocretin) circuits to pathologies of motivation. Br J Pharmacol 2024; 181(22): 4430–4449.

24. Calipari ES, Espana RA. Hypocretin/orexin regulation of dopamine signaling: implications for reward and reinforcement mechanisms. Front Behav Neurosci 2012; 6: 54.

25. James MH, Aston-Jones G. Orexin Reserve: A Mechanistic Framework for the Role of Orexins (Hypocretins) in Addiction. Biol Psychiatry 2022; 92(11): 836–844.

26. Aston-Jones G, Smith RJ, Moorman DE, Richardson KA. Role of lateral hypothalamic orexin neurons in reward processing and addiction. Neuropharmacology 2009; 56 Suppl 1(Suppl 1): 112–121.

27. Brodnik ZD, Alonso IP, Xu W, Zhang Y, Kortagere S, Espana RA. Hypocretin receptor 1 involvement in cocaine-associated behavior: Therapeutic potential and novel mechanistic insights. Brain Res 2020; 1731: 145894.

28. Kallo I, Omrani A, Meye FJ, de Jong H, Liposits Z, Adan RAH. Characterization of orexin input to dopamine neurons of the ventral tegmental area projecting to the medial prefrontal cortex and shell of nucleus accumbens. Brain Struct Funct 2022; 227(3): 1083–1098.

29. Schultz W, Dayan P, Montague PR. A neural substrate of prediction and reward. Science 1997; 275(5306): 1593–1599.

30. Flagel SB, Clark JJ, Robinson TE, Mayo L, Czuj A, Willuhn I, Akers CA, Clinton SM, Phillips PE, Akil H. A selective role for dopamine in stimulus-reward learning. Nature 2011; 469(7328): 53–57.

31. Berke JD. What does dopamine mean? Nat Neurosci 2018; 21(6): 787–793.

32. Korotkova TM, Sergeeva OA, Eriksson KS, Haas HL, Brown RE. Excitation of ventral tegmental area dopaminergic and nondopaminergic neurons by orexins/hypocretins. J Neurosci 2003; 23(1): 7–11.

33. Nakamura T, Uramura K, Nambu T, Yada T, Goto K, Yanagisawa M, Sakurai T. Orexin-induced hyperlocomotion and stereotypy are mediated by the dopaminergic system. Brain Res 2000; 873(1): 181–187.

34. Borgland SL, Taha SA, Sarti F, Fields HL, Bonci A. Orexin A in the VTA is critical for the induction of synaptic plasticity and behavioral sensitization to cocaine. Neuron 2006; 49(4): 589–601.

35. Borgland SL, Storm E, Bonci A. Orexin B/hypocretin 2 increases glutamatergic transmission to ventral tegmental area neurons. Eur J Neurosci 2008; 28(8): 1545–1556.

36. Thomas CS, Mohammadkhani A, Rana M, Qiao M, Baimel C, Borgland SL. Optogenetic stimulation of lateral hypothalamic orexin/dynorphin inputs in the ventral tegmental area potentiates mesolimbic dopamine neurotransmission and promotes reward-seeking behaviours. Neuropsychopharmacology 2022; 47(3): 728–740.

37. Bonifazi A, Del Bello F, Giorgioni G, Piergentili A, Saab E, Botticelli L, Cifani C, Micioni Di Bonaventura E, Micioni Di Bonaventura MV, Quaglia W. Targeting orexin receptors: Recent advances in the development of subtype selective or dual ligands for the treatment of neuropsychiatric disorders. Med Res Rev 2023; 43(5): 1607–1667.

38. Fenwick EM, Marty A, Neher E. A patch-clamp study of bovine chromaffin cells and of their sensitivity to acetylcholine. J Physiol 1982; 331: 577–597.

39. Baimel C, Lau BK, Qiao M, Borgland SL. Projection-Target-Defined Effects of Orexin and Dynorphin on VTA Dopamine Neurons. Cell Reports 2017; 18(6): 1346–1355.

40. Asahi S, Egashira S, Matsuda M, Iwaasa H, Kanatani A, Ohkubo M, Ihara M, Morishima H. Development of an orexin-2 receptor selective agonist, [Ala(11), D-Leu(15)]orexin-B. Bioorg Med Chem Lett 2003; 13(1): 111–113.

41. Tsuneoka Y, Funato H. Whole Brain Mapping of Orexin Receptor mRNA Expression Visualized by Branched In Situ Hybridization Chain Reaction. eNeuro 2024; 11(2).

42. Simon P, Dupuis R, Costentin J. Thigmotaxis as an index of anxiety in mice. Influence of dopaminergic transmissions. Behav Brain Res 1994; 61(1): 59–64.

43. File SE, Mabbutt PS, Hitchcott PK. Characterisation of the phenomenon of “one-trial tolerance” to the anxiolytic effect of chlordiazepoxide in the elevated plus-maze. Psychopharmacology (Berl) 1990; 102(1): 98–101.

44. Treit D, Menard J, Royan C. Anxiogenic stimuli in the elevated plus-maze. Pharmacol Biochem Behav 1993; 44(2): 463–469.

45. Zhou H, Yu CL, Wang LP, Yang YX, Mao RR, Zhou QX, Xu L. NMDA and D1 receptors are involved in one-trial tolerance to the anxiolytic-like effects of diazepam in the elevated plus maze test in rats. Pharmacol Biochem Behav 2015; 135: 40–45.

46. Blomeley C, Garau C, Burdakov D. Accumbal D2 cells orchestrate innate risk-avoidance according to orexin signals. Nat Neurosci 2017.

47. Frussa-Filho R, Ribeiro Rde A. One-trial tolerance to the effects of chlordiazepoxide in the elevated plus-maze is not due to acquisition of a phobic avoidance of open arms during initial exposure. Life Sci 2002; 71(5): 519–525.

48. File SE, Zangrossi H, Jr., Viana M, Graeff FG. Trial 2 in the elevated plus-maze: a different form of fear? Psychopharmacology (Berl) 1993; 111(4): 491–494.

49. Bari A, Dalley JW, Robbins TW. The application of the 5-choice serial reaction time task for the assessment of visual attentional processes and impulse control in rats. Nat Protoc 2008; 3(5): 759–767.

50. Bariselli S, Glangetas C, Tzanoulinou S, Bellone C. Ventral tegmental area subcircuits process rewarding and aversive experiences. J Neurochem 2016; 139(6): 1071–1080.

51. Solie C, Girard B, Righetti B, Tapparel M, Bellone C. VTA dopamine neuron activity encodes social interaction and promotes reinforcement learning through social prediction error. Nat Neurosci 2022; 25(1): 86–97.

52. Gunaydin LA, Deisseroth K. Dopaminergic Dynamics Contributing to Social Behavior. Cold Spring Harb Symp Quant Biol 2014; 79: 221–227.

53. Dawson M, Terstege DJ, Jamani N, Tsutsui M, Pavlov D, Bugescu R, Epp JR, Leinninger GM, Sargin D. Hypocretin/orexin neurons encode social discrimination and exhibit a sex-dependent necessity for social interaction. Cell Rep 2023; 42(7): 112815.

54. Abbas MG, Shoji H, Soya S, Hondo M, Miyakawa T, Sakurai T. Comprehensive Behavioral Analysis of Male Ox1r (-/-) Mice Showed Implication of Orexin Receptor-1 in Mood, Anxiety, and Social Behavior. Front Behav Neurosci 2015; 9: 324.

55. Moy SS, Nadler JJ, Perez A, Barbaro RP, Johns JM, Magnuson TR, Piven J, Crawley JN. Sociability and preference for social novelty in five inbred strains: an approach to assess autistic-like behavior in mice. Genes Brain Behav 2004; 3(5): 287–302.

56. Cryan JF, Holmes A. The ascent of mouse: advances in modelling human depression and anxiety. Nat Rev Drug Discov 2005; 4(9): 775–790.

57. McGraw M, Christensen C, Nelson H, Li AJ, Qualls-Creekmore E. Divergent changes in social stress-induced motivation in male and female mice. Physiol Behav 2025; 291: 114787.

58. Figueiredo Cerqueira MM, Castro MML, Vieira AA, Kurosawa JAA, Amaral Junior FLD, Siqueira Mendes FCC, Sosthenes MCK. Comparative analysis between Open Field and Elevated Plus Maze tests as a method for evaluating anxiety-like behavior in mice. Heliyon 2023; 9(4): e14522.

59. Xiao X, Yeghiazaryan G, Cremer AL, Backes H, Kloppenburg P, Hausen AC. Deficiency of Orexin Receptor Type 1 in Dopaminergic Neurons Increases Novelty-Induced Locomotion and Exploration. 2023.

60. Ekstrand MI, Terzioglu M, Galter D, Zhu S, Hofstetter C, Lindqvist E, Thams S, Bergstrand A, Hansson FS, Trifunovic A, Hoffer B, Cullheim S, Mohammed AH, Olson L, Larsson NG. Progressive parkinsonism in mice with respiratory-chain-deficient dopamine neurons. Proc Natl Acad Sci U S A 2007; 104(4): 1325–1330.

61. Backman CM, Malik N, Zhang Y, Shan L, Grinberg A, Hoffer BJ, Westphal H, Tomac AC. Characterization of a mouse strain expressing Cre recombinase from the 3’ untranslated region of the dopamine transporter locus. Genesis 2006; 44(8): 383–390.

62. Giros B, Jaber M, Jones SR, Wightman RM, Caron MG. Hyperlocomotion and indifference to cocaine and amphetamine in mice lacking the dopamine transporter. Nature 1996; 379(6566): 606–612.

63. Costa KM, Schenkel D, Roeper J. Sex-dependent alterations in behavior, drug responses and dopamine transporter expression in heterozygous DAT-Cre mice. Sci Rep 2021; 11(1): 3334.

64. Li S, Franken P, Vassalli A. Bidirectional and context-dependent changes in theta and gamma oscillatory brain activity in noradrenergic cell-specific Hypocretin/Orexin receptor 1-KO mice. Scientific Reports 2018; 8(1): 15474.

65. Zalocusky KA, Ramakrishnan C, Lerner TN, Davidson TJ, Knutson B, Deisseroth K. Nucleus accumbens D2R cells signal prior outcomes and control risky decision-making. Nature 2016; 531(7596): 642–646.

66. van den Pol AN, Gao XB, Obrietan K, Kilduff TS, Belousov AB. Presynaptic and postsynaptic actions and modulation of neuroendocrine neurons by a new hypothalamic peptide, Hypocretin/Orexin. Journal of Neuroscience 1998; 18(19): 7962–7971.

67. Patyal R, Woo EY, Borgland SL. Local hypocretin-1 modulates terminal dopamine concentration in the nucleus accumbens shell. Front Behav Neurosci 2012; 6: 82.

68. Lambe EK, Aghajanian GK. Hypocretin (orexin) induces calcium transients in single spines postsynaptic to identified thalamocortical boutons in prefrontal slice. Neuron 2003; 40(1): 139–150.

69. Aracri P, Banfi D, Pasini ME, Amadeo A, Becchetti A. Hypocretin (orexin) regulates glutamate input to fast-spiking interneurons in layer V of the Fr2 region of the murine prefrontal cortex. Cereb Cortex 2015; 25(5): 1330–1347.

70. Kroener S, Chandler LJ, Phillips PE, Seamans JK. Dopamine modulates persistent synaptic activity and enhances the signal-to-noise ratio in the prefrontal cortex. PLoS One 2009; 4(8): e6507.

71. Ott T, Nieder A. Dopamine and Cognitive Control in Prefrontal Cortex. Trends Cogn Sci 2019; 23(3): 213–234.

72. Stalter M, Westendorff S, Nieder A. Dopamine Gates Visual Signals in Monkey Prefrontal Cortex Neurons. Cell Rep 2020; 30(1): 164–172 e164.

73. Salamone JD, Correa M. The Neurobiology of Activational Aspects of Motivation: Exertion of Effort, Effort-Based Decision Making, and the Role of Dopamine. Annu Rev Psychol 2024; 75: 1–32.

74. Andolina D, Puglisi-Allegra S, Ventura R. Strain-dependent differences in corticolimbic processing of aversive or rewarding stimuli. Front Syst Neurosci 2014; 8: 207.

75. Freet CS, Arndt A, Grigson PS. Compared with DBA/2J mice, C57BL/6J mice demonstrate greater preference for saccharin and less avoidance of a cocaine-paired saccharin cue. Behav Neurosci 2013; 127(3): 474–484.

76. Zweifel LS, Fadok JP, Argilli E, Garelick MG, Jones GL, Dickerson TM, Allen JM, Mizumori SJ, Bonci A, Palmiter RD. Activation of dopamine neurons is critical for aversive conditioning and prevention of generalized anxiety. Nat Neurosci 2011; 14(5): 620–626.

77. Stelly CE, Haug GC, Fonzi KM, Garcia MA, Tritley SC, Magnon AP, Ramos MAP, Wanat MJ. Pattern of dopamine signaling during aversive events predicts active avoidance learning. Proc Natl Acad Sci U S A 2019; 116(27): 13641–13650.

78. Zeidler Z, Gomez MF, Gupta TA, Shari M, Wilke SA, DeNardo LA. Prefrontal dopamine activity is critical for rapid threat avoidance learning. bioRxiv 2025.

79. Blouin AM, Fried I, Wilson CL, Staba RJ, Behnke EJ, Lam HA, Maidment NT, Karlsson KAE, Lapierre JL, Siegel JM. Human hypocretin and melanin-concentrating hormone levels are linked to emotion and social interaction. Nat Commun 2013; 4: 1547.

80. Crawley JN. Twenty years of discoveries emerging from mouse models of autism. Neurosci Biobehav Rev 2023; 146: 105053.

81. Crawley JN. Designing mouse behavioral tasks relevant to autistic-like behaviors. Ment Retard Dev Disabil Res Rev 2004; 10(4): 248–258.

82. Silverman JL, Thurm A, Ethridge SB, Soller MM, Petkova SP, Abel T, Bauman MD, Brodkin ES, Harony-Nicolas H, Wohr M, Halladay A. Reconsidering animal models used to study autism spectrum disorder: Current state and optimizing future. Genes Brain Behav 2022; 21(5): e12803.

83. Genovese A, Butler MG. The Autism Spectrum: Behavioral, Psychiatric and Genetic Associations. Genes (Basel) 2023; 14(3).

84. Sato M, Nakai N, Fujima S, Choe KY, Takumi T. Social circuits and their dysfunction in autism spectrum disorder. Mol Psychiatry 2023; 28(8): 3194–3206.

85. Muroya S, Funahashi H, Yamanaka A, Kohno D, Uramura K, Nambu T, Shibahara M, Kuramochi M, Takigawa M, Yanagisawa M, Sakurai T, Shioda S, Yada T. Orexins (hypocretins) directly interact with neuropeptide Y, POMC and glucose-responsive neurons to regulate Ca 2+ signaling in a reciprocal manner to leptin: orexigenic neuronal pathways in the mediobasal hypothalamus. Eur J Neurosci 2004; 19(6): 1524–1534.

86. Ma X, Zubcevic L, Bruning JC, Ashcroft FM, Burdakov D. Electrical inhibition of identified anorexigenic POMC neurons by orexin/hypocretin. J Neurosci 2007; 27(7): 1529–1533.

87. Belle MD, Hughes AT, Bechtold DA, Cunningham P, Pierucci M, Burdakov D, Piggins HD. Acute suppressive and long-term phase modulation actions of orexin on the mammalian circadian clock. J Neurosci 2014; 34(10): 3607–3621.

88. Izawa S, Fusca D, Jiang H, Heilinger C, Hausen AC, Wunderlich FT, Steuernagel L, Kloppenburg P, Bruning JC. Orexin/hypocretin receptor 2 signaling in MCH neurons regulates REM sleep and insulin sensitivity. Cell Rep 2025; 44(2): 115277.

89. Kukkonen JP, Turunen PM. Cellular Signaling Mechanisms of Hypocretin/Orexin. Front Neurol Neurosci 2021; 45: 91–102.

90. Hatcher-Solis C, Fribourg M, Spyridaki K, Younkin J, Ellaithy A, Xiang G, Liapakis G, Gonzalez-Maeso J, Zhang H, Cui M, Logothetis DE. G protein-coupled receptor signaling to Kir channels in Xenopus oocytes. Curr Pharm Biotechnol 2014; 15(10): 987–995.

91. Ishibashi M, Gumenchuk I, Miyazaki K, Inoue T, Ross WN, Leonard CS. Hypocretin/Orexin Peptides Alter Spike Encoding by Serotonergic Dorsal Raphe Neurons through Two Distinct Mechanisms That Increase the Late Afterhyperpolarization. J Neurosci 2016; 36(39): 10097–10115.

92. Brodie MS, McElvain MA, Bunney EB, Appel SB. Pharmacological reduction of small conductance calcium-activated potassium current (SK) potentiates the excitatory effect of ethanol on ventral tegmental area dopamine neurons. J Pharmacol Exp Ther 1999; 290(1): 325–333.

93. Um KB, Kwak S, Cheon SH, Kim J, Hwang SK. AST-001 Improves Social Deficits and Restores Dopamine Neuron Activity in a Mouse Model of Autism. Biomedicines 2023; 11(12).

94. Stott JB, Greenwood IA. G protein betagamma regulation of KCNQ-encoded voltage-dependent K channels. Front Physiol 2024; 15: 1382904.

95. Morris LS, Costi S, Hameed S, Collins KA, Stern ER, Chowdhury A, Morel C, Salas R, Iosifescu DV, Han MH, Mathew SJ, Murrough JW. Effects of KCNQ potassium channel modulation on ventral tegmental area activity and connectivity in individuals with depression and anhedonia. Mol Psychiatry 2025.

96. Korotkova TM, Eriksson KS, Haas HL, Brown RE. Selective excitation of GABAergic neurons in the substantia nigra of the rat by orexin/hypocretin in vitro. Regul Pept 2002; 104(1-3): 83–89.

97. Liu C, Xue Y, Liu MF, Wang Y, Liu ZR, Diao HL, Chen L. Orexins increase the firing activity of nigral dopaminergic neurons and participate in motor control in rats. J Neurochem 2018; 147(3): 380–394.

98. Bian K, Liu C, Wang Y, Xue Y, Chen L. Orexin-B exerts excitatory effects on nigral dopaminergic neurons and alleviates motor disorders in MPTP parkinsonian mice. Neurosci Lett 2021; 765: 136291.

99. Leger L, Sapin E, Goutagny R, Peyron C, Salvert D, Fort P, Luppi PH. Dopaminergic neurons expressing Fos during waking and paradoxical sleep in the rat. J Chem Neuroanat 2010; 39(4): 262–271.

100. Cho JR, Treweek JB, Robinson JE, Xiao C, Bremner LR, Greenbaum A, Gradinaru V. Dorsal Raphe Dopamine Neurons Modulate Arousal and Promote Wakefulness by Salient Stimuli. Neuron 2017; 94(6): 1205–1219 e1208.

101. Korchynska S, Rebernik P, Pende M, Boi L, Alpar A, Tasan R, Becker K, Balueva K, Saghafi S, Wulff P, Horvath TL, Fisone G, Dodt HU, Hokfelt T, Harkany T, Romanov RA. A hypothalamic dopamine locus for psychostimulant-induced hyperlocomotion in mice. Nat Commun 2022; 13(1): 5944.

102. Lu X, Xue J, Lai Y, Tang X. Heterogeneity of mesencephalic dopaminergic neurons: From molecular classifications, electrophysiological properties to functional connectivity. FASEB J 2024; 38(3): e23465.

103. Ogawa Y, Kanda T, Vogt K, Yanagisawa M. Anatomical and electrophysiological development of the hypothalamic orexin neurons from embryos to neonates. J Comp Neurol 2017; 525(18): 3809–3820.

104. Grafe LA, Cornfeld A, Luz S, Valentino R, Bhatnagar S. Orexins Mediate Sex Differences in the Stress Response and in Cognitive Flexibility. Biol Psychiatry 2017; 81(8): 683–692.

105. Grafe LA, Bhatnagar S. The contribution of orexins to sex differences in the stress response. Brain Res 2020; 1731: 145893.

106. Xiao X, Yeghiazaryan G, Eggersmann F, Cremer AL, Backes H, Kloppenburg P, Hausen AC. Deficiency of orexin receptor type 1 in dopaminergic neurons increases novelty-induced locomotion and exploration. Elife 2025; 12.

107. Summers CH, Yaeger JDW, Staton CD, Arendt DH, Summers TR. Orexin/hypocretin receptor modulation of anxiolytic and antidepressive responses during social stress and decision-making: Potential for therapy. Brain Research 2020; 1731.

108. Jacobson LH, Hoyer D, de Lecea L. Hypocretins (orexins): The ultimate translational neuropeptides. J Intern Med 2022; 291(5): 533–556.

109. McGregor R, Wu M-F, Thannickal T, Siegel J. Opiate anticipation, opiate induced anatomical changes in hypocretin (Hcrt, orexin) neurons and opiate induced microglial activation are blocked by the dual Hcrt receptor antagonist suvorexant, while opiate analgesia is maintained. bioRxiv 2024.

110. Chaki S. Orexin receptors: possible therapeutic targets for psychiatric disorders. Psychopharmacology (Berl) 2025.

111. Zhuang C, Cao Y, Lu J, Zhou Y, Liu Y, Li Y. The orexin/hypocretin system in dementia-related neurological disorders: a double-edged sword in cognitive impairment. Psychopharmacology (Berl) 2025.

112. Williams JT, Bolli MH, Brotschi C, Sifferlen T, Steiner MA, Treiber A, Gatfield J, Boss C. Discovery of Nivasorexant (ACT-539313): The First Selective Orexin-1 Receptor Antagonist (SO1RA) Investigated in Clinical Trials. J Med Chem 2024; 67(4): 2337–2348.

113. Reid MS, Tafti M, Nishino S, Sampathkumaran R, Siegel JM, Mignot E. Local administration of dopaminergic drugs into the ventral tegmental area modulates cataplexy in the narcoleptic canine. Brain Res 1996; 733(1): 83–100.

114. Dauvilliers Y, Mignot E, Del Rio Villegas R, Du Y, Hanson E, Inoue Y, Kadali H, Koundourakis E, Meyer S, Rogers R, Scammell TE, Sheikh SI, Swick T, Szakacs Z, von Rosenstiel P, Wu J, Zeitz H, Murthy NV, Plazzi G, von Hehn C. Oral Orexin Receptor 2 Agonist in Narcolepsy Type 1. N Engl J Med 2023; 389(4): 309–321.

115. Dauvilliers Y, Plazzi G, Mignot E, Lammers GJ, Del Rio Villegas R, Khatami R, Taniguchi M, Abraham A, Hang Y, Kadali H, Lamberton M, Sheikh S, Stukalin E, Neuwirth R, Swick TJ, Tanaka S, von Hehn C, von Rosenstiel P, Wang H, Cai A, Naylor M, Olsson T. Oveporexton, an Oral Orexin Receptor 2-Selective Agonist, in Narcolepsy Type 1. N Engl J Med 2025; 392(19): 1905–1916.

116. Markovic T, Pedersen CE, Massaly N, Vachez YM, Ruyle B, Murphy CA, Abiraman K, Shin JH, Garcia JJ, Yoon HJ, Alvarez VA, Bruchas MR, Creed MC, Morón JA. Pain induces adaptations in ventral tegmental area dopamine neurons to drive anhedonia-like behavior. Nature Neuroscience 2021; 24(11): 1601–1613.

## Supplemental References

117. Meneghini S, Modena D, Colombo G, Coatti A, Milani N, Madaschi L, Amadeo A, Becchetti A. The beta2(V287L) nicotinic subunit linked to sleep-related epilepsy differently affects fast-spiking and regular spiking somatostatin-expressing neurons in murine prefrontal cortex. Prog Neurobiol 2022; 214: 102279.

118. Franklin KBJ, Paxinos G. Paxinos and Franklin’s The mouse brain in stereotaxic coordinates. Fourth edition. edn, 1 volume (unpaged)pp.

119. Zhang HY, Gao M, Shen H, Bi GH, Yang HJ, Liu QR, Wu J, Gardner EL, Bonci A, Xi ZX. Expression of functional cannabinoid CB(2) receptor in VTA dopamine neurons in rats. Addict Biol 2017; 22(3): 752–765.

120. Khaliq ZM, Bean BP. Pacemaking in dopaminergic ventral tegmental area neurons: depolarizing drive from background and voltage-dependent sodium conductances. J Neurosci 2010; 30(21): 7401–7413.

121. Ford CP. The role of D2-autoreceptors in regulating dopamine neuron activity and transmission. Neuroscience 2014; 282: 13–22.

122. Margolis EB, Toy B, Himmels P, Morales M, Fields HL. Identification of rat ventral tegmental area GABAergic neurons. PLoS One 2012; 7(7): e42365.

123. Tzanoulinou S, Riccio O, de Boer MW, Sandi C. Peripubertal stress-induced behavioral changes are associated with altered expression of genes involved in excitation and inhibition in the amygdala. Transl Psychiatry 2014; 4(7): e410.

124. Walf AA, Frye CA. The use of the elevated plus maze as an assay of anxiety-related behavior in rodents. Nat Protoc 2007; 2(2): 322–328.

125. Contestabile A, Casarotto G, Girard B, Tzanoulinou S, Bellone C. Deconstructing the contribution of sensory cues in social approach. Eur J Neurosci 2021; 53(9): 3199–3211.

126. Tzanoulinou S, Musardo S, Contestabile A, Bariselli S, Casarotto G, Magrinelli E, Jiang YH, Jabaudon D, Bellone C. Inhibition of Trpv4 rescues circuit and social deficits unmasked by acute inflammatory response in a Shank3 mouse model of Autism. Mol Psychiatry 2022; 27(4): 2080–2094.

127. Sasamori H, Ohmura Y, Kubo T, Yoshida T, Yoshioka M. Assessment of impulsivity in adolescent mice: A new training procedure for a 3-choice serial reaction time task. Behav Brain Res 2018; 343: 61–70.

128. Asinof SK, Paine TA. The 5-choice serial reaction time task: a task of attention and impulse control for rodents. J Vis Exp 2014; (90): e51574.

129. Tsutsui-Kimura I, Ohmura Y, Izumi T, Yamaguchi T, Yoshida T, Yoshioka M. The effects of serotonin and/or noradrenaline reuptake inhibitors on impulsive-like action assessed by the three-choice serial reaction time task: a simple and valid model of impulsive action using rats. Behav Pharmacol 2009; 20(5-6): 474–483.

130. Humby T, Wilkinson L, Dawson G. Assaying aspects of attention and impulse control in mice using the 5-choice serial reaction time task. Curr Protoc Neurosci 2005; **Chapter 8:** Unit 8 5H.

131. Cain CK. Avoidance Problems Reconsidered. Curr Opin Behav Sci 2019; 26: 9–17.

132. Tabibnia G. An affective neuroscience model of boosting resilience in adults. Neurosci Biobehav Rev 2020;115: 321–350.

133. Diehl MM, Bravo-Rivera C, Quirk GJ. The study of active avoidance: A platform for discussion. Neurosci Biobehav Rev 2019; 107: 229–237.

